# An atypical class of non-coding small RNAs produced in rice leaves upon bacterial infection

**DOI:** 10.1101/2021.03.05.432875

**Authors:** Ganna Reshetnyak, Jonathan M. Jacobs, Florence Auguy, Coline Sciallano, Lisa Claude, Clemence Medina, Alvaro L. Perez-Quintero, Aurore Comte, Emilie Thomas, Adam Bogdanove, Ralf Koebnik, Boris Szurek, Anne Dievart, Christophe Brugidou, Severine Lacombe, Sebastien Cunnac

**Affiliations:** PHIM Plant Health Institute, Univ Montpellier, IRD, CIRAD, INRAE, Institut Agro, Montpellier, France; Department of Plant Pathology, The Ohio State University, Columbus, OH, United States, USA 43201 and Infectious Disease Institute, The Ohio State University, Columbus, OH, United States, USA 43201; Department of Agricultural Biology, Colorado State University, Fort Collins, CO, USA; Plant Pathology and Plant-Microbe Biology Section, School of Integrative Plant Science, Cornell University, Ithaca, NY, United States; AGAP, Univ Montpellier, CIRAD, INRAE, Institut Agro, Montpellier, France

**Keywords:** small RNA, *Xanthomonas oryzae*, rice, type III secretion, disease, plant

## Abstract

Non-coding small RNAs (sRNA) act as mediators of gene silencing and regulate plant growth, development and stress responses. Early insights into plant sRNAs established a role in antiviral defense and they are now extensively studied across plant-microbe interactions. Here, sRNA sequencing discovered a class of sRNA in rice (*Oryza sativa*) specifically associated with foliar diseases caused by *Xanthomonas oryzae* bacteria. *Xanthomonas*-induced small RNAs (xisRNAs) loci were distinctively upregulated in response to diverse virulent strains at an early stage of infection producing a single duplex of 20-22nt sRNAs. xisRNAs production was dependent on the Type III secretion system, a major bacterial virulence factor for host colonization. xisRNA loci overlap with annotated transcripts sequences often encoding protein kinase domain proteins. A number of the corresponding rice *cis-*genes have documented functions in immune signaling and some xisRNA loci coincide with the coding sequence of a conserved kinase motif. xisRNAs exhibit features of small interfering RNAs and their biosynthesis depend on canonical components *OsDCL1* and *OsHEN1.* xisRNA induction possibly mediates post-transcriptional gene silencing but they do not broadly suppress *cis*-genes expression on the basis of mRNA-seq data. Overall, our results identify a group of unusual sRNAs with a potential role in plant-microbe interactions.

## INTRODUCTION

RNA silencing represents a fundamental mechanism of gene regulation in eukaryotes and orchestrates a wide range of basic cellular and organismal processes. In plants, RNA silencing coordinates major biological functions such as maintenance of genome integrity and inheritance, development and environmental responses (1, 2). Non-coding small RNAs (sRNAs) determine the specificity of RNA silencing. sRNAs are typically 20-24 nucleotides (nt) in size and derive from a double stranded RNA (dsRNA) precursor that is diced into a single or multiple distinct sRNA duplexes with 2-3nt overhangs by Dicer-like proteins (DCL). In plants, the released sRNAs duplexes are universally methylated on the 2′-OH of the 3′ terminal ribose by HEN1. This methyl group protects them from rapid uridylation and subsequent degradation (3). The mature sRNA duplex ‘guide’ strand is selectively loaded into an ARGONAUTE (AGO) protein component of RNA-Induced gene Silencing Complexes to mediate either transcriptional gene silencing (TGS) or post-transcriptional gene silencing (PTGS) according to nucleic acids sequence complementarity between the sRNAs and their target RNA (4). TGS typically depends on 24nt small interfering RNAs (siRNAs) and involves DNA methylation and chromatin reprogramming. PTGS, on the other hand, guided by 21nt micro and siRNAs, leads to mRNA cleavage and mRNA translational inhibition (1).

Several major classes of endogenous plant regulatory sRNAs have been defined based on the specifics of their biogenesis pathways and their mode of action (3, 5): (i) micro RNAs (miRNAs) are arguably the best characterized class of sRNAs. Hairpins of *MIRNA* gene transcripts (pri-miRNA) are cleaved and diced by the plant DCL1 protein to generate a single miRNA duplex. The guide strand of a miRNA duplex typically associates with AGO1 to exert PTGS. (ii) other hairpin small interfering RNAs (hp-siRNAs) also derive from a single-stranded RNA with inverted repeats that folds into a hairpin structure but whose processing does not follow the precise pattern associated with miRNAs. The remaining three main classes of sRNAs pertain to siRNAs whose dsRNA precursor is made up of two complementary RNA molecules and include (iii) natural antisense transcript (NAT) siRNAs, (iv) secondary siRNAs and (v) heterochromatic siRNAs. ∼24nt heterochromatic sRNAs typically mediate TGS. NAT-siRNAs are produced from two independently transcribed but complementary precursors (Natural Antisense Transcripts or NATs) and can directly target one molecule of their precursor (*cis*-) or target mRNAs that do not derive from their locus of origin (*trans*-). Secondary siRNAs biogenesis is triggered by an upstream sRNA through an RNA-dependent RNA polymerase (RDR) catalyzed reaction and often results in the amplification of the original sRNA signal both in terms of quantity and complexity. In this category, phased secondary siRNAs (phasiRNAs) and *trans*-acting siRNAs are generated following the conversion of a single-stranded miRNA cleavage product to a double stranded precursor by RDR6 (6).

The role of sRNAs in plant biotic interactions is increasingly being appreciated and while the defense against invading RNA viruses was one of the earliest functions ascribed to plant RNA silencing, it is well-established that multiple sRNA pathways are also critical for plant immune responses to bacterial, fungal or oomycete pathogens (7). In nature, healthy plants rely on their innate immune system to perceive and fight off pathogens. Their first line of defense is the detection of conserved molecular signatures, such as bacterial flagella or fungal chitin. Plasma membrane localized receptor-kinases and receptor-like proteins detect these Pathogen-Associated Molecular Patterns (PAMPs). PAMPs perception triggers phosphorylation cascades that ultimately signal the deployment of defense mechanisms to hinder pathogen invasion. This process is referred to as PAMP-triggered immunity (PTI) (8). Plant receptor-kinases (a.k.a. RLKs for Receptor-Like Kinases) have a monophyletic origin, are related to the *Drosophila* Pelle kinases and mammalian INTERLEUKIN1 RECEPTOR-ASSOCIATED KINASE (IRAK), and have been classified into the RLK/Pelle kinase family (9, 10). This family also includes Receptor-Like Cytoplasmic Kinases (RLCKs) that lack the transmembrane and extracellular domains and function in intracellular signal transduction as substrates of receptor-kinase complexes (11). Several plant miRNAs whose expression is modulated in response to PAMPs were shown to be important to efficiently combat pathogen attacks downstream of the perception of PAMP signals (7). Adapted pathogens have evolved mechanisms to counteract eukaryotic immune responses. For instance, many Gram-negative bacterial pathogens employ a Type Three Secretion System (T3SS) to inject effector proteins directly into eukaryotic host cells to modulate their physiology and suppress defenses (12). Similar to Viral Suppressors of RNA silencing plant-pathogenic bacteria and oomycetes have been shown to use their effector repertoire to dampen sRNA silencing activity and compromise plant immunity (13). As a counter-defense mechanism, plants evolved resistance (R) proteins that recognize pathogen virulence factors through a mechanism called Effector-Triggered Immunity (ETI) which often restricts pathogen growth and infection by triggering programmed cell death (14). The role of sRNA as positive mediators of immunity is not limited to the PTI layer, and both TGS and PTGS also regulate the ETI layer (7).

Rice (*Oryza sativa*) is a major crop in global agriculture and stands as an economic and nutritional cornerstone in the developing world. Rice possesses RNA silencing pathways that are both analogous to *A. thaliana* and idiosyncratic (15). It is noteworthy that multigene families of core RNA silencing components seem to have undergone significant expansion in rice (8 DCLs, 19 AGOs and 5 RDRs) (16). Global rice production is limited by plant diseases caused by pathogens such as viruses, fungi, nematodes and bacteria. *Xanthomonas oryzae* pv. *oryzae* (*Xoo*) and *X. oryzae* pv. *oryzicola* (*Xoc*) are the pathogenic agents that cause major diseases of rice: bacterial leaf blight (BLB) and bacterial leaf streak (BLS), respectively. *Xoo* causes yield losses of 20-30% that culminate at 50% in some regions of Asia. *Xoo* and *Xoc* form distinct phylogenetic groups that are further delimited as a function of their geographic origin. Even though they belong to the same *X. oryzae* (*Xo*) species, they have distinct lifestyles *in planta*. *Xoo* colonizes the xylem, a nutrient-limiting, minimal plant environment used for water, mineral and organic acid transport; meanwhile *Xoc* enters through stomata and infects the plant mesophyll to cause leaf streak (17). *Xoo* and *Xoc* employ a suite of virulence factors, including the T3SS, that quantitatively contribute to disease development. Mutants lacking the T3SS are unable to translocate effector proteins into host cells and are non-virulent. These bacteria possess over 32 effectors, including a unique class of Transcription Activator-Like Effector (TALE) that is particularly large in *Xoo* and *Xoc*. TALEs are important T3SS-delivered effectors that activate protein-encoding gene expression and reprogram the transcriptome of the host’s cells by mimicking eukaryotic transcription factors (12).

Recently, the discovery of trans-kingdom RNAi or *trans*-species small RNA silencing, has shed a new light on the role of sRNAs as mobile effectors modulating the outcome of plant biotic interactions (7, 18). For example, the necrotrophic fungal pathogen *Botrytis cinerea* transfers its own sRNA to silence defense genes of its host plants Arabidopsis and tomato. Pathogen effectors acting as RNA silencing suppressors and pathogen *trans*-species sRNAs are both delivered into the plant cell to prevent defense. However, while the former proteinaceous effectors repress silencing, the latter ribonucleic effectors instead, exploit it. There is still limited evidence in support of a third virulence strategy where proteinaceous effectors would leverage RNA silencing to promote immune suppression and susceptibility. Recently however, a *Xoo* T3SS effector-induced rice miRNA was shown to exert PTGS on a *R* gene conferring quantitative resistance to BLB (*19*).

With a few exceptions, such as *Arabidopsis* AtlsiRNA-1 corresponding to a novel class of 30–40 nt long siRNAs and nat-siRNAATGB2 that contribute to the ETI launched after recognition of a bacterial effector (7), the available functional data on the role of endogenous plant sRNAs in regulating the disease process has focused almost exclusively on miRNAs and their role in promoting immunity. For example, in rice, the expression of multiple miRNAs changes in response to the blast fungus *Magnaporthe oryzae* (20) and *Xoo* (21, 22). Here, we aimed at examining all sRNAs associated with the disease process at a genome-wide scale and used an unbiased small RNA deep-sequencing data analysis approach on rice inoculated with *Xo* strains. Unusual 20-22nt small RNAs were found to be generated from annotated protein coding loci in the *Xo* infected leaf in a T3SS-dependant manner and required the archetypal sRNA biogenesis pathways components *OsDCL1* and *OsHEN1*. These sRNAs often mapped to *cis*-genes previously implicated in immune signaling, but did not appear to generally operate through PTGS of their *cis*-gene. This study provides insight into an intriguing group of sRNAs that may participate in gene silencing for plant disease development.

## MATERIAL AND METHODS

### Bacterial strains and plant inoculation

The *Xo* strains used in this study are described in Supplementary Table S4. The isolates were maintained in 15% glycerol at -80°C. All *Xoo* strains were cultivated on PSA (10 g/liter peptone, 10 g/liter sucrose, 1 g/liter glutamic acid, 15 g/liter Bacto Agar), supplemented with kanamycin (50 μg/g/ ml), rifampicin (100 μg/g/ml) and/or tetracycline (5 μg/g/ml) when needed for the propagation of mutant and complemented strains. For plant infiltration, frozen bacterial stocks were spread on Petri dishes with PSA and incubated at 28°C. On the morning of the experiment, bacterial suspensions were prepared in distilled water and injected with a 1ml needleless syringe on the abaxial face of the youngest fully expended leaf of 4 weeks old rice plants. Bacterial suspensions were adjusted at OD_600nm_=0.5 except for infiltration of *waf1-2* homozygous plants that were found in preliminary experiments to require a three fold more concentrated suspension in order to deliver an equivalent number of bacteria in the leaf mesophyll at t_0_. After infiltration, plants were kept in the same greenhouse growth conditions until sample collection.

### Plant material and growth conditions

Plants were grown under greenhouse conditions with cycles of 12 h of light at 28°C and 80% relative humidity and 12 h of dark at 25°C and 70% relative humidity. The following rice genotypes were used: cultivar Nipponbare (23), the DCL1-IR transgenic line #43-6 in the Nipponbare background (24), Kinmaze variety and *waf1-2* mutant derivate (25) and a heterozygous *dcl4-1* line (26). For PCR genotyping with primers listed in Supplementary Table S5 and the GoTaq G2 DNA Polymerase (Promega) according to manufacturer’s instructions, template rice genomic DNA was extracted from leaf samples using a standard MATAB-based protocol (27).

### RNA extraction

Unless stated otherwise, ∼2cm fragments were collected from a leaf of three independently infiltrated plants at the specified time point after inoculation and pooled in 2 mL microtubes containing 3 steel beads and immediately frozen in liquid nitrogen. For routine northern blot and Q- RT-PCR experiment, total RNA was extracted using 1ml of Tri reagent (MRC) following the manufacturer’s instructions and resuspended in nuclease-free water. For Illumina sequencing, the total RNA was treated with the TURBO DNA-free Kit (ThermoFisher) following instructions from the manufacturer and further purified with the mirVana Isolation Kit (ThermoFisher).

### RNA Next-Generation Sequencing

The sRNA-seq data for the BAI3 dataset was generated by the Fasteris company (Geneva, Switzerland). Briefly, sRNA libraries were prepared using the Illumina TruSeq small RNA kit with polyacrylamide gel purified 18-25nt sRNA fractions and sequenced on a portion of a Illumina HiSeq 2000 lane, 1x50 single-reads. The sRNA-seq data for the Diversity dataset was generated by the Novogene company (Hong Kong, China) using a NEBNext Small RNA Library Prep Set for Illumina kit optimized by Novogene on the supplied total RNAs. Ultimately, size selected 18-47bp inserts were sequenced on a Illumina HiSeq (1x50bp). The ‘BAI3’ mRNA stranded sequencing was performed on a HiSeq 2000 lane (2x100bp) by the Fasteris company (Geneva, Switzerland) with libraries prepared following a dir-mRNA-dUTP protocol that includes poly-adenylated transcripts selection followed by cDNA library construction using Illumina TruSeq Stranded mRNA Library Prep kit. Demultiplexing of sequencing reads was performed by the sequence providers.

### Northern Blots and Quantative RT-PCR

Low molecular weight northen blots were performed using previously described standard procedures (28). Briefly, 10μg/g of total RNA in volume of 40μg/L in 1X gel loading buffer II (Ambion) were loaded on a on a 17,5% denaturing acrylamide gel and electrophoresed at 80V, transferred to a Hybond NX membrane (Amersham) and crosslinked with UV. The membrane was then pre-hybridized 30 min with PerfectHyb buffer (Sigma-Aldrich). The radioactively labeled probe was first prepared with γ[32 P]ATP ATP and DNA oligonucleotides listed in Supplementary Table S5 with a T4 polynucleotide kinase kit (Promega), purified on MicroSpin G-25 (GE Healthcare) column and denatured 2 min at 95°C. The RNAs on the membrane were hybridized with the probe overnight at 40°C in PerfectHyb buffer and washed three times with a 2X SSC 0.1% SDS solution for 5min. Radioactive signals were imaged using a Typhoon phosphorimager (Amersham). Contrast adjustments were applied to blot hybridized with individual probes in a linear fashion and faithfully represent original signal information. Before hybridization with a new probe, northern membranes were stripped with three 10min washes in near-boiling 0.1% SDS and re-equilibrated in 2X SSC. For RT-qPCR, after contaminant DNA removal with the TURBO DNA-free Kit (ThermoFisher), 1μg/g RNA was reverse transcribed into cDNA using SuperScriptIII (ThermoFisher) and random hexamers as specified by the manufacturer. The reactions were diluted 5 times in water before quantitiative real-based protocol time PCR on a LightCycler LC480 (Roche), using SYBR-based protocol based Mesa Blue qPCR Mastermix (Eurogentec) and the oligonucleotides primers listed in Supplementary Table S5. Transcript levels in a sample were calculated using the ΔΔCt method from two technical replicate reactions from the same cDNA preparation using the *OsEF*-11α gene for normalization.

### Deep sequencing data primary analysis

The first stage of our sRNA sequencing data processing pipeline included a 3’ adaptor trimming step performed with AdapterRemoval (v. 2.2.2 - https://github.com/MikkelSchubert/adapterremoval) using these key parameters : “--mm 5 --minadapteroverlap 3 --minlength 18 --maxlength 50”. Illumina reads were mapped onto the nuclear rice genome with bwa aln (29) version 0.7.17 with default parameters and the resulting files were further processed with samtools version 1.7 (http://www.ncbi.nlm.nih.gov/pubmed/19505943). All downstream analysis steps were conducted separately for each individual dataset. sRNA loci inference was performed with ShortStack (v. 3.8.5) (30) using the following key parameters : “--dicermin 20 --dicermax 24 --mincov 2rpm --pad 75 --foldsize 400 --strand_cutoff 0.8”. Read counts were obtained using the summarizeOverlaps function from the GenomicAlignments Bioconductor package. Experimental sRNA loci that accumulated less than 30 total reads across all libraries in the dataset were excluded. Likewise, sRNA loci that overlapped with rice loci with feature types “tRNA”, “rRNA” or “pseudogenic_tRNA” in the NCBI gff annotation file for GCF_001433935.1_IRGSP-1.0 (ftp://ftp.ncbi.nlm.nih.gov/genomes/all/GCF/001/433/935/GCF_001433935.1_IRGSP-1.0/GCF_001433935.1_IRGSP-1.0_genomic.gff.gz) were excluded from differential expression (DE) analysis.

For replicated datasets, standard DE analysis was performed with the DESeq2 R Bioconductor package (31). For normalization, size factors were estimated using the “iterate” method and the “shorth” function to compute a location for a sample. Dispersions were fitted using the “parametric”, by default, or the “local” method, in case estimates did not converge. Two-tailed Wald test were used for DE tests where the alternative hypotheses was that the absolute value of the log2 transform of the fold change ratio is greater than 2. Adjusted p-values were computed with the Benjamini & Hochberg method.

For unreplicated datasets, DE analysis used the DESeq package (32) with default parameters except that a “blind” value for the method parameter and a value “fit-only” for the sharingMode parameter were applied instead in the calls to the estimateDispersions function. As above, dispersions were fitted using the “parametric”, by default, or the “local” method, in case estimates did not converge. Benjamini & Hochberg adjusted p-values were computed for the null hypothesis that the value of the log2 transform of the fold change ratio is equal to 0. The latest version of our primary sRNA processing workflow was written into a Snakemake pipeline and is available on gitHub (https://github.com/Aucomte/sRNAmake).

For the analysis of differential gene expression using mRNA-seq data, we applied the ARMOR pipeline (33) with the following parameter values specified in the config file: additional_salmon_index: “-k 31”, additional_salmon_quant: “--seqBias –gcBias”. Note that whenever available in the sample metadata information, a ‘experiment’ factor term was included in the statistical model for sRNA loci and gene DE analysis in order to block for experimental batch effects.

### Secondary bioinformatic analysis

Rice Nipponbare genomic data including genome sequence, gene annotation and transcripts sequences were downloaded from the MSU website (http://rice.plantbiology.msu.edu/pub/data/Eukaryotic_Projects/o_sativa/annotation_dbs/pseudomolecules/version_7.0/all.dir/). MIRNA annotation was obtained from MirBase (v22) (34). Unless otherwise stated, pri-miRNAs loci considered in these analysis correspond to those reference rice sRNA loci annotations in the Plant small RNA gene annotations database (35) that are flagged as “MIRNA” (“type” field), that are also listed in miRBase (“source” field has value “miRBase21”) and for which an overlapping experimental sRNA loci has been identified in all the datasets considered in this study.

We performed Singular Enrichment Analysis for GO Slim Gene ontology terms in the “Rice MSU7.0 nonTE gene” dataset on the AgriGO website (http://bioinfo.cau.edu.cn/agriGO/ - date: 10/09/2020) with the list of *cis*-gene identifiers and default parameter values (Fisher statistical test, Yekutieli p-value correction).

To build the rice and Arabidopsis kinase phylogenetic tree available in Supplementary File S1, the hmmsearch program (36) was used to search for Hidden Markov Model (HMM) kinase profile (PF00560) into the rice MSU7 and Arabidopsis TAIR10 (http://www.arabidopsis.org/) proteomes. Kinase domain sequences were aligned using MAFFT (37). This alignment was cleaned with TrimAl configured to remove sites with more than 80% of gaps (38). Based on this alignment, a phylogenetic tree has been computed with the Fasttree method (approximately-maximum-likelihood method) (39).

The search for domain hits in the InterPro database (40) used InterProScan linux executable and dataset (v. 5.41-78.0) as well as the Panther data (v. 14.1) with default parameters on the set of MSU7 annotated protein with an underlying xisRNA duplex region fully included within the coding sequence.

The search for major xisRNA reads similarities in bacterial genomes used the blastn tool of the BLAST+ suite (v. 2.10.1) with these relevant parameters: “-task \”blastn-short\” -max_target_seqs 5-perc_identity 80”. The blast database was created with the following strain and NCBI assembly accessions: BAI3 (GCA_003031385.1), MAI1 (GCA_003031365.1), BLS256 (GCA_000168315.3), BAI11 (GCA_000940825.1).

The secondary bioinformatic and statistical analysis that generated the graphics and results included in this study have been essentially conducted using the R Statistical programming language and packages, notably from the Bioconductor project (41). The corresponding code and output files have been deposited in a dataverse with the doi:10.23708/RXIXM3 DOI.

## RESULTS

### Small RNA sequencing of rice leaves infected with Xoo BAI3 identifies sRNA loci that are upregulated in a T3SS-dependant manner

To understand the molecular mechanisms underlying plant-bacterial interactions, we defined host sRNA species that may be involved in the disease process. To gain genome-wide insight, we generated sRNA Illumina sequencing data from RNA collected from the Japonica *O. sativa* rice cultivar Nipponbare leaves confronted by pathogenic Xoo. Briefly, sRNAs were analyzed from leaf samples taken over two biological replicates 24 hours post-infiltration (hpi) with the wild-type African Xoo reference strain BAI3, a non -virulent T3SS mutant, which is unable to inject virulence effectors (herein referred to as BAI3H) or a water control (H2O). Following sRNA reads mapping, we performed genome segmentation based on sRNA reads coverage profile along genomic sequences with the ShortStack script (30) to infer potential sRNA generating genomic loci on the Nipponbare genome (23), and tested for differential accumulation of sRNA reads in treatment comparisons as summarized in Supplementary Figure S1. From a list of 27,096 experimental sRNA loci, our differential accumulation pipeline identified those with a log2-transformed fold-change ratio across treatment comparisons equal or above 2 and an associated adjusted p-value equal or above 0.05. With these stringent criteria, 341 sRNAs loci were found to be differentially expressed (DE) in at least one of the three treatment comparisons tested (Supplementary Table S1). Regardless of the comparison, all DE sRNA loci were upregulated in tissues inoculated with bacteria (either BAI3 or BAI3H) compared to mock inoculated (H2O) leaves. Because the T3SS is an essential component of infection, we aimed to define sRNA loci that were induced in a T3SS- dependent fashion. We required that <u>X</u>anthomonas-induced <u>s</u>mall RNA (xisRNA) loci be significantly induced in both the BAI3 vs H2O and the BAI3 vs BAI3H comparisons but not in the BAI3H vs H20 comparison where no T3SS effect is expected to occur. To ensure that xisRNA loci would not correspond to repetitive sequences and would be enriched for potential regulatory non-coding sRNAs, we further required that ≥ 75% of the reads at these loci mapped uniquely in the genome and that ≥ 50% of the reads had a size between 20 and 24 nt. Applying these criteria to the set of 341 DE sRNAs loci resulted in a total of 64 xisRNA loci.

Thus, NGS of sRNAs identified a number of potential regulatory rice sRNA loci where mapping reads predominantly accumulate in the context of a T3SS-dependent *Xoo* BAI3 infection.

### BAI3 xisRNAs overlap with exons of annotated protein-coding genes and have features of siRNAs

To characterize xisRNA loci and notably evaluate whether they may generate regulatory non-coding sRNAs, we first analyzed the distribution of xisRNA loci length (Figure 1A) and found that with a median of 106 base pairs (bp), their length is significantly different from the length (median = 449bp) of other experimentally defined sRNA loci (Two-sided Wilcoxon rank sum test: W = 1547014, p = 1.02e-27). In addition, inspecting the distribution of the proportion of reads mapping to the top genomic strand or ‘strandedness ratio’ in experimental sRNA loci (Figure 1B), revealed that, as expected, a vast majority of the experimental sRNA loci overlapping MirBase pri-miRNA genomic intervals that are also annotated in the Plant small RNA gene annotations database (35) have strandedness ratios of 0 or 1. In contrast, the bulk of xisRNA and other experimental sRNA loci have values that vary inside this interval and are composed of reads partitions that map to either strands of their cognate genomic locus, reminiscent of siRNA loci. xisRNA loci expression in non-inducing conditions is extremely low or even null (Figure 1C and D). In contrast, upon BAI3 infection, xisRNA loci reads are rather abundant. Even if overall lower, their normalized expression is not at odd with pri-miRNA loci levels (Figure 1D). Importantly, the expression level of the most highly expressed loci, namely, xisRNA023, 01, 02, 05, 07, 09 and 10 reaches several thousand rpm in leaves inoculated with virulent BAI3 bacteria (Figure 1C). Regarding read sizes distributions, no major global bias in *MIRNA* genes-derived reads sizes and coverage patterns were detected (Supplementary Figure S2). While this suggests that read populations in our dataset are not anomalously biased, read length distribution for xisRNA loci was broader than for *MIRNA* genes (Figure 1C). Furthermore, with a few exceptions (e.g. xiRNA059, 62, 12, 53 and 51), xisRNA read sizes ranged overall from 20 to 22nt, which is comparable to rice siRNAs.

**Figure 1:**
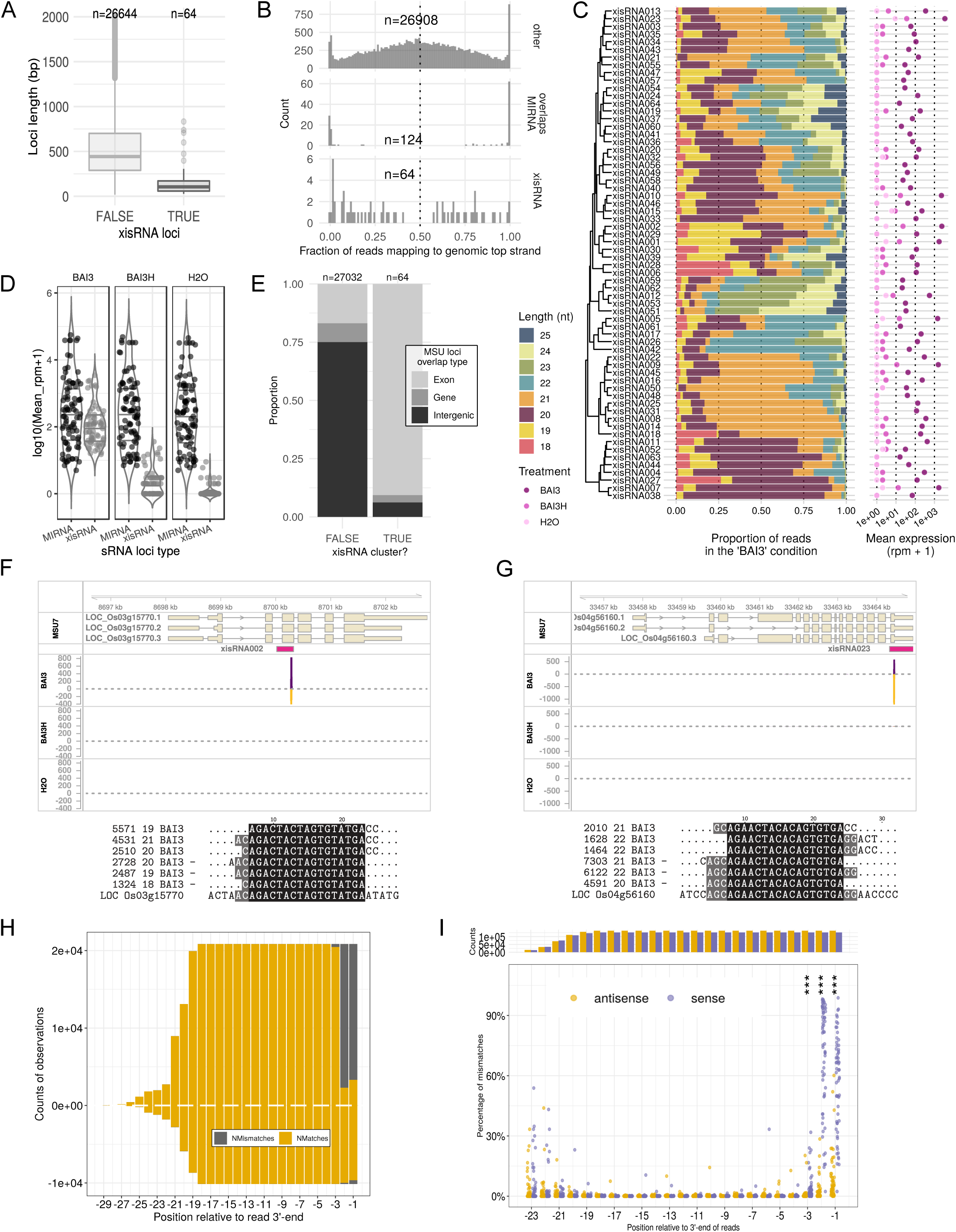
Xanthomonas-induced small RNAs (xisRNA) signatures and loci possess distinctive attributes. **A.** Box plot representation (median, 1^st^ and 3^rd^ quartile and 1.5 * IQR) of the distribution of the length of the experimental sRNA loci, contrasted as a function of whether they correspond to a xisRNA or not. Note that 388 loci were excluded from the plot because their lenght is above 2000bp. **B.** Distribution of the fraction of reads mapping to the genomic top strand for experimental sRNA loci contrasted as a function of whether the loci correspond to xisRNA, MIRNA or other loci. **C.** Distribution of sRNA reads length and mean expression at the 64 xisRNA loci. The hierarchical clustering of xisRNA loci is based on the profiles of reads proportions for individual length values. Mean expression values in each treatment (shades of pink) are represented on a log10 scale. **D.** Distribution of mean normalized expression in reads per million (rpm) for pri-miRNA loci (n=87) versus xisRNA loci (n=64) in libraries derived from various leaf inoculation treatments. **E.** Proportions of the type of rice annotation features overlapping with experimental sRNA loci, contrasted as a function of whether sRNA loci correspond to a xisRNA or not. Overlaps are included in the counts if spanning at least 20bp. Experimental sRNA loci overlapping both a gene and an exon annotation range are classified as ‘Exon’. Those overlapping a gene annotation element only are classified as ‘Gene’. Others are classified as ‘Intergenic’. **F.** Genome browser snapshot of the MSU7 LOC_Os03g15770 locus annotation overlapping with the xisRNA002 ShortStack segment and read coverage tracks (reads per million) for the various experimental treatments. Shorter exon boxes correspond to untranslated regions. Coverage of reads mapping to the top genomic strand is represented with positive values and a violet colored area whereas coverage of reads mapping to the opposite strand is represented with negative values and a golden colored area. The multiple sequence alignment below the genome browser graph corresponds to the three most abundant unique mapping signatures per strand of the ShortStack loci as well as the genomic sequence (LOC ID). In the sequence title, the two numbers indicate, respectively, the count of reads with this sequence in the “BAI3” treatment libraries and the length of the unique sequence. The “-” sign indicates that the original read sequences map to the complementary strand of the genomic locus. The displayed sequences are thus the reverse complement of the original read sequences. **G.** Same as E but for the LOC_Os04g56160 locus region overlapping with xisRNA023. **H.** Counts of nucleotides matching (Nmatches) or not matching (Nmismatches) the xisRNA002 loci genomic sequences along the read alignment region. Positions on the x-axis are expressed relative to the sRNA reads 3’-end (last nucleotide is -1). Positive values are computed for reads mapping to the annotated sense strand of the LOC_Os03g15770 locus. Conversely, negative values correspond to reads mapping to the antisense strand of the annotated loci. **I.** Distribution of genomic sequence mismatch ratios along relative positions within reads for xisRNA loci. Each data point represents the ratio of the number of mismatches over the total number of comparisons with the genomic sequence for that position of all the reads mapping to a xisRNA locus in the sense orientation (violet) or antisense orientation (golden) relative to the MSU annotated transcript overlapping with this xisRNA locus. Positions where a two-sided Wilcoxon signed rank test measuring differences in xisRNA mismatch ratios between read orientations were significant (BY adjusted p-value ≤ 0.05) are marked with a “*” sign. Here, three “*” indicate an adj. P-val. ≤ 0.001. The bar graph on the top reports on the total count of comparisons for each orientation.

In order to relate xisRNA loci with existing rice miRNA annotation, we asked whether they overlap with *O. sativa MIRNA* gene loci annotated in MirBase but found no match. Consistent with this observation, none of the xisRNA loci was classified by ShortStack as a possible MIRNA locus (category ‘N14’, ‘N15’ or ‘Y’), suggesting that their features are incompatible with their being previously undocumented MIRNA genes. We also examined the overlap of experimental sRNA loci with annotated genes and transcripts in the Rice Genome Annotation Project (MSU7 annotation) (23). As illustrated in Figure 1E, the distribution of the overlap type with MSU loci is significantly different for xisRNA loci as compared to other experimental sRNA loci (Two-sided Fisher’s Exact test p-value = 6.05e-38). Notably, 58 out of 64 xisRNA loci (91%) overlap with an exon of an annotated MSU loci which is significantly greater than the proportion obtained for other experimental sRNA loci (One-sided Fisher’s Exact test adjusted p-value = 1.57e-37).

In summary, xisRNA loci are generally short and do not correspond to documented or putative *MIRNA* genes but a majority of them overlap with exons of annotated rice genes. Analogous to siRNAs, these loci are composed of 20-22 nucleotides-long reads mapping to both strands of the genomic sequence and that can also accumulate to substantial levels.

### BAI3 xisRNAs loci presumably generate single sRNA duplexes

A majority of xisRNA loci overlap with exons of annotated protein-coding genes. We therefore examined the coverage profile of mapped xisRNAs signatures along the Nipponbare genome sequence in regions encompassing annotated MSU loci (see coverage plots in our dataverse). For example, highly expressed and typical xisRNA002 and xisRNA023 loci display a ∼25bp-wide coverage peak enclosed in an annotated exon in the BAI3 library track alone (Figures 1F and G). These T3SS-dependant peaks comprise reads mapping in either sense or antisense orientation. To extent this observation, we retrieved the three most abundant unique read signatures for each strand of the xisRNA loci as well as the sequence of the annotated MSU locus transcript in this window and created the multiple alignments reproduced underneath the genome plots of Figure 1F and 1G. One striking feature of these alignments is that xisRNA major reads sizes and the pattern of complementarity between sense and anti-sense reads with short (∼2nt) 3’-overhangs are evocative of guide-passenger strands of sRNAs duplexes resulting from DCL processing. Another discernible feature is that in contrast to major reads in the antisense orientation relative to the annotated *cis*-transcript, the last 2-3 3’-end nucleotides of major reads in the sense orientation do not match the transcript sequence. To evaluate whether the presence of untemplated 3’-most nucleotides is a general feature of xisRNA sense reads, we computed the counts of matches and mismatches relative to the genomic sequence for individual positions along read sequences in each xisRNA loci and contrasted them as a function of read orientation (exemplified for xisRNA002 in Figure 1H). This further enabled to derive the position- and strand-specific mismatch ratios distributions plotted in Figure 1I for all xisRNA loci. Consistent with our preliminary observation based on major xsiRNA signatures for xisRNA002 and xisRNA023, statistical tests for differences in xisRNA mismatch ratios between read orientations for individual positions were significant (adjusted p-value ≤ 0.05) only for the last first, second and third nucleotides (positions -1, -2 and -3) where xisRNA loci displayed overall higher proportions of mismatches for sense reads than for antisense reads. The substitution matrix of genomic sequence versus reads sequences for positions -2 and -1 in Supplementary Figure S3A indicates that xisRNA sense reads untemplated 3’ nucleotides are overwhelmingly biased for cytosines, independently of the nature of the underlying genomic nucleotide. To rule out the possibility that this predominance of 3’ untemplated nucleotides in sense xsiRNA reads would be a global artifact of our Illumina sequencing experiment, we conducted the same analysis for a subset of non-xisRNA experimental sRNA loci and did not conclude on any significant differences in mismatch ratios (Supplementary Figure S3B). In contrast, testing differences in sense reads mismatch ratios between pri-miRNA transcripts and xisRNA loci indicated that these ratios were significantly higher for xisRNA than for pri-miRNA loci Supplementary Figure S3C).

In short, we obtained evidence that while similar to non-coding small RNAs loci, individual xisRNA loci exhibit unconventional features in that they are essentially composed of complementary reads reminiscent of a single sRNA duplex with those reads in the sense orientation relative to an annotated *cis*-mRNA having a high proportion of untemplated cytosines in the last 2-3 3’ nucleotides.

### **X.** *oryzae* strains from multiple sublineages induce xisRNAs

In order to independently validate the assumption that xisRNA NGS reads correspond to genuine sRNA species, we performed standard low molecular weight northern blots with probes designed to detect the moderately to highly expressed xisRNA duplex molecules that are complementary to annotated cis-transcripts. In agreement with this view and as shown in Figure 2A and Supplementary Figure S4, for all tested xisRNA loci, following Nipponbare leaf infiltration with the BAI3 strain, a T3SS-dependent signal was detected no earlier that 24hpi and gradually increased in intensity up to 72hpi. This corresponds to the latest sampled time point (Supplementary Figure S4) because severe disease symptoms complicated RNA extraction for later time points.

**Figure 2:**
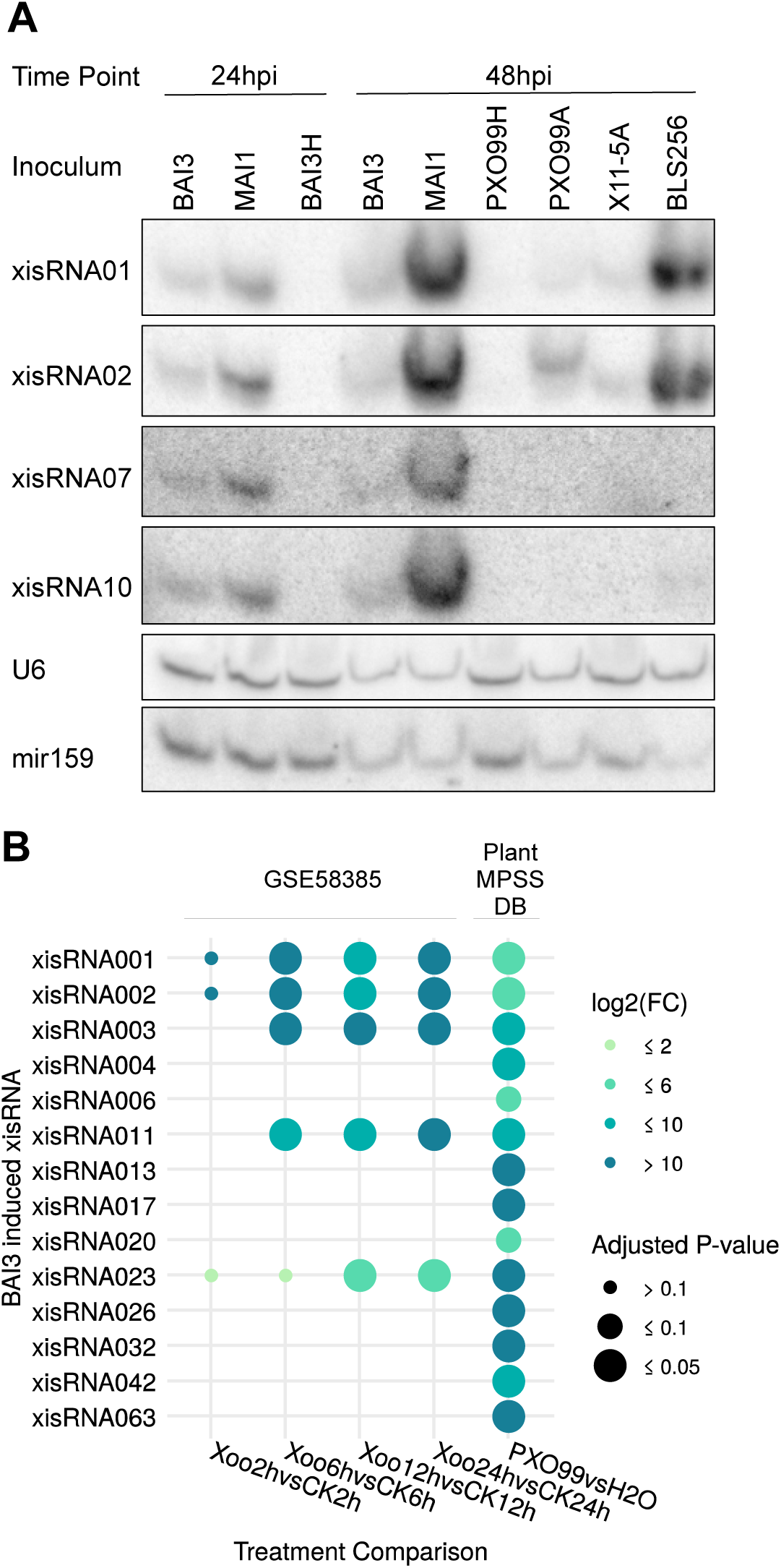
A subset of xisRNAs are detected in northern blots and exhibit contrasted expression patterns in response to phylegentically diverse Xo strains. **A.** Autoradiographs obtained after RNA gel northern blot analysis conducted with specific oligonucleotide probes hybridized to total RNA extracted from Nipponbare leaves infiltrated with the indicated strains and collected at the indicated time points (hpi = hours post inoculation). The xisRNA probes detect RNAs complementary to the sense strand of the xisRNA *cis*-gene. The BAI3H and PXO99H strains correspond to a T3SS defective mutant derivative of the BAI3 and PXO99A virulent strains, respectively. Individual images derive from successive stripping, hybridization and detection rounds of the same membrane. Equivalent results were obtained in two other similar experiments. **B.** Differential expression statistics of xisRNA loci found to be induced by the Asian Xoo PXO99 strain in public sRNA-seq datasets (time course experiment after leaf cliping [GSE58385]ATP or Plant MPSS database). The treatment comparisons (bottom labels) involve tissue samples inoculated with PXO99 versus a mock control. The plot displays xisRNA loci with a log2(fold change) ≥ 1 and an BH adjusted p-values ≤ 0.1 in at least one comparison.

The broad genetic diversity of the *X. oryzae* species is structured as a function of the continent of origin of the strains and their parasitic strategy as captured under the concept of pathovar (*Xoo* or *Xoc*). We were curious to determine if the capacity to trigger xisRNA expression was specific to *Xoo* BAI3 alone or if it is a conserved feature of the phylogenetically diverse *Xo* species. Northern blot experiments were conducted with RNA samples extracted from leaves infiltrated with reference strains representative of the main genetic and pathotypic groups: African (BAI3, MAI1), Asian (PXO99A, KACC 10331) or American (X11-5A) *Xoo* strains as well as African (e.g. MAI10) or Asian *Xoc* strains (e.g. BLS256). As shown in Figure 2A and Supplementary Figure S5, *X. oryzae* strains were broadly capable of eliciting xisRNA accumulation with contrasting strength and specificities. In qualitative terms, African *Xoo* and *Xoc* strains induced all tested xisRNAs with the exception of xisRNA007 which seems to be specific to African *Xoo*. The Asian *Xoo* strains and the American *Xoo* strain were less active and fainter but reproducible signals could be detected essentially for xisRNA001 and xisRNA002. In the case of PXO99A, these signals were also T3SS-dependent. While xisRNA023 was not tested for the US *Xo* strain X11-5A, xisRNA023 signals appeared to be specific to *Xoo* pathovar strains as these sRNAs were readily detected in response to both Asian and African *Xoo* strains (Supplementary Figure S5).

As a complementary way to establish that xisRNA induction is not limited to African *Xoo* infection, we exploited the dataset generated by Zhao *et al.* (22) in the NCBI GEO database (accession GSE58385) where sRNAs expression temporal dynamics in *Xoo* PXO99 versus mock-inoculated rice leaves was profiled at early time points (2, 6, 12 and 24hpi) following leaf clipping inoculation. A second dataset deposited by B. Meyers and B. Yang and including Nipponbare sRNA sequencing data 24hpi with the Azacytidine-resistant derivative of PXO99 strain, PXO99A or mock was downloaded from the Plant MPSS databases (42). These independent libraries were used as input in our genome segmentation and DE pipeline with modifications to account for a lack of replicates (see Material and Methods). The DE statistics of experimental sRNA loci whose expression changed in relevant treatment comparisons and that overlapped with BAI3 xisRNA loci are summarized in Figure 2B. Several of these experimental sRNA loci were also found to be up- regulated by PXO99 relative to water controls (eg xisRNA01, 02, 23). Coverage profiles of sRNA reads in xisRNA loci further indicate that PXO99-induced reads map exactly to the same location as BAI3-induced reads (see coverage plots in our dataverse). In addition to confirming that PXO99 also induces xisRNA accumulation, the higher sensitivity of sRNA-seq applied to leaf clipping assays samples further indicates that some xsiRNAs accumulates as early as six hours following inoculation in an assay that more closely recapitulates a natural infection of xylem vessels as opposed to massive infiltration of the leaf mesophyll.

The broad xisRNA induction capacity in the genetic diversity of both *Xo* pathovars emphasizes the relevance of these sRNAs as evolutionary conserved molecular markers of early rice infection events and implies a potential functional role in this process.

### A broader xisRNAs inventory

Having established that xisRNA accumulation is a widespread molecular feature of rice infection by *Xo* pathogens raised the question of whether other strains induced novel xisRNA loci that were not captured by our initial BAI3-specific sRNA-seq experiment. To address this, we performed a second set of rice sRNA-seq experiments with three biological replicates per treatment using the same methodological and analytical framework as above. African *Xoo* MAI1 strain was selected because in northern blot experiments it strongly induced xisRNAs. As representatives of the *Xoc* pathovar, the African BAI11 strain (genome sequenced in (43)) and the Asian BLS256, together with a T3SS mutant derivative (BLS256H), were also selected to investigate T3SS-dependency and *Xoc* pathovar specific xisRNAs. The setup of these experiments as well as the outcome of the primary genome segmentation and DE analysis are summarized in Supplementary Figure S6 and Supplementary Table S2. In order to identify rice xisRNA loci in this dataset designated as the ‘Diversity’ dataset, we applied the same criteria as for the first ‘BAI3’ dataset. Regarding the MAI1 and BAI11 strains for which no T3SS minus mutant derivative was included, only loci induced in the “MAI1vsH2O” or the “BAI11vsH2O” but not in “BLS256HvsH2O” comparison were defined as xisRNAs loci. Across both datasets, a total of 163 distinct loci met the criteria defined for xisRNA loci, 99 of which were not identified in the BAI3 dataset. Supplementary Table S3 records expression measures, features of mapping sRNA reads and annotated *cis*-genes information for these loci while Figure 3 depicts a graphical representation of this data for the 60 most highly expressed of these loci. The clustering of DE patterns across comparisons (Figure 3 and Supplementary Figure S7A) is complex but pointed to few main classes of xisRNA loci: 16 are broadly induced by all tested *Xo* strains (e.g. xisRNA001, 02), 22 are induced by African *Xoo* and *Xoc* strains (e.g. xisRNA009, 22) and 57 are specifically induced by the Asian Xoc BLS256.

**Figure 3:**
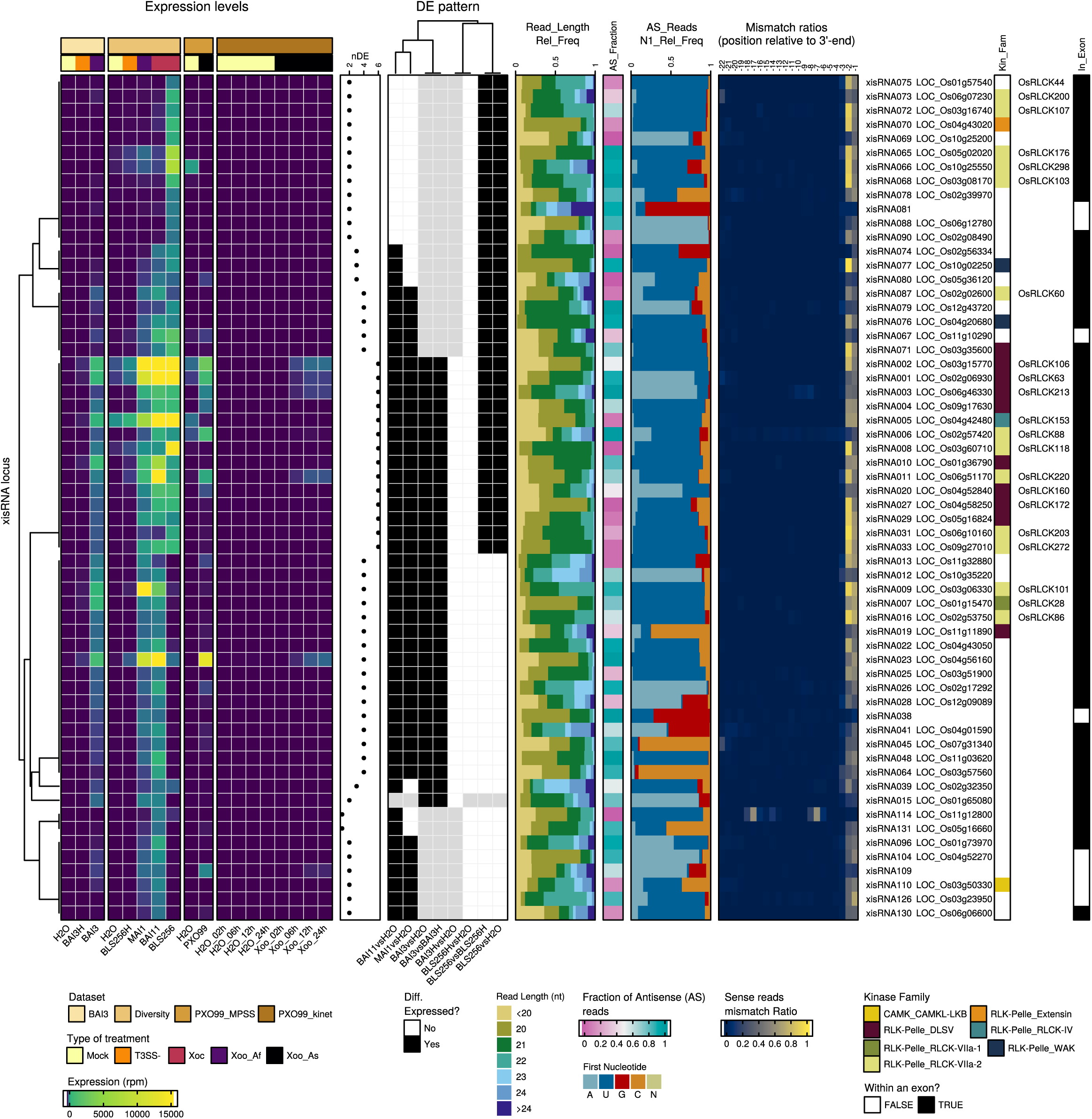
A summary of xisRNAs features in the analyzed sRNA-seq datasets. In the complex heatmap layout, the top 60 xisRNA loci with the highest maximum treatment mean expression across datasets are described row-wise. The first component, on the left, reports on expression (reads per million) in the various inoculation treatments of the different datasets. In the type of treatment legend, the “Xoo_Af” and “Xoo_As” labels stand for African *Xoo* and Asian *Xoo* strains, respectively. The DE patterns component shows whether xisRNA loci were detected as DE in various comparisons. The “nDE” dot plot depicts the number of comparisons where the xisRNA locus was deferentially expressed. Cells in gray indicate that the xisRNA locus was not considered in the dataset because the loci was not detected by ShortStack or because reads in this dataset did not pass our xisRNA loci features criteria. The subsequent plots on the right depicts the size distribution of reads (“Read_Length_Rel_Freq”), the fraction of antisense reads (“AS_Fraction”), the proportion of antisense reads with distinct 5’- end first nucleotide (“AS_Reads_N1_Rel_Freq”) and the specific nucleotide mismatch ratios for relative positions -22 to -1 of antisense reads using xisRNA loci mapping reads from the “BAI3” and “Diversity” datasets. The MSU7 identifier of the gene overlapping with a xisRNA duplex locus is indicated together with the kinase subfamilly into which the encoded protein is classified (“Kin_Fam.” column) as well as the RLCK protein name, when relevant and available. Finally, the “In_Exon” column indicates whether the xisRNA duplex loci is fully included in an exon of an annotated mRNA of the gene. All numerical values reported in the graphic were computed using reads mapping to the xisRNA duplex loci. Note that the antisense polarity is defined relative to the orientation of the underlying *cis*-gene. To compute values that incorporate a notion of polarity for xisRNA loci that do not overlap with an annotated gene, reads mapping to the top strand were arbitrarily considered to be in the sense orientation.

In order to ascertain again that the sRNA reads mapping at ShortStack-inferred dataset-specific xisRNA loci in this extended inventory were congruent with the hypothesis that xisRNAs derive from a duplex of sRNAs molecules, we inspected the coverage of all reads from both datasets. With the exception of xisRNA094 for which the shape of read coverage is not consistent with a sRNA duplex model, all other loci displayed a single major ∼25bp wide coverage peak (see coverage plot in our dataverse). We therefore defined these shorter 25bp genomic intervals as ‘xisRNA duplex loci’ (see xisRNA loci-specific genome browser views in our dataverse) . As illustrated in Figure 3 and exhaustively recorded in Supplementary Table S3, xisRNA duplex loci reads in the BAI3 and Diversity datasets exhibit the same distinguishing features (strandedness ratios, read size distribution and 3’ untemplated nucleotides) as described before. Individual AGO family members preferentially select guide sRNAs with a specific 5′ nucleotide (4). We therefore computed the composition of the 5’ nucleotide of xisRNA duplex loci reads as a function of their polarity to examine potential bias. As shown in Supplementary Figure 7B and Figure 3, duplex loci reads complementary to the *cis*-gene strand have a proportion of 5’-T that is well above the proportion of Ts in the rice genome. A hallmark of sRNA loaded into rice AGO1s complexes is a strong sequence bias for 5’ Uracil (44). This implies that the composition of *cis*-gene antisense xisRNA 5’-nucleotide is in line with their possible loading into an AGO1 complex.

In summary, the joint analysis of our sRNA-seq datasets generated a larger collection of xisRNA loci detected independently with diverse *Xo* strains and confirmed that the main features of xisRNAs described in the BAI3 dataset were also conserved in the Diversity dataset. Furthermore, cis-gene antisense xisRNAs might be sorted into OsAGO1 complexes to silence *cis-*genes.

### Many protein products of loci overlapping with xisRNAs harbor kinase domains and some are involved in immune responses signaling

Of the 163 xisRNA duplex loci, 91% (149/163) overlapped with an MSU annotated gene (*cis*-gene) and 75% (122/163) were included within an annotated exon. To examine if those genes had related functions, we performed Plant GO Slim term enrichment analysis on the AgriGO website with the list of *cis*-gene identifiers (Supplementary Figure S8). In the Molecular Function ontology domain, we noticed a significant enrichment (FDR ≤ 0.05) of the “RNA binding” term. Among several predicted proteins with RNA-binding domains (e.g. RNA recognition motif, KH domain [IPR004087]ATP, Tudor-SN [IPR016685]ATP, DEAD-box RNA helicase), the presence of two genes overlapping with African *Xo* strains specific xisRNAs and coding for products involved in sRNA silencing pathways was especially noteworthy: the ARGONAUTE family member *OsAGO13* (16) associated with xisRNA064 and the DICER-LIKE family member *OsDCL4*/*SHO1* (26, 45) associated with xisRNA022 (Figure 3, Supplementary Table S3).

In the Molecular Function GO category, the most significantly enriched terms are connected to “receptor activity” and “kinase activity”, with no less than 64 xisRNA loci overlapping genes (46%) annotated with this latter term. In order to gain some insight on the family composition of kinase domain containing products of rice xisRNA loci *cis*-genes, they were annotated in Figure 3 and Supplementary Table S3 based on the kinase classification of Lehti-Shiu and Shiu (9) and the rice OsRLCK gene family classification of Vij et al. (46). Consistent with GO enrichment analysis, the proportions of the different kinase groups in the xisRNA set is different from the whole genome composition (two-sided Fisher’s Exact test p-value=5e-04) and a one-sided post-hoc Fisher’s exact test indicated that 5 subgroups (Extensin, WAK, DLSV, RLCK-IV, RLCK-VIIa-2) of the RLK/Pelle protein kinase family are significantly enriched in the xisRNA *cis*-gene products list (Holm method adjusted p-values < 0.05). RLK/Pelle subfamilies such as DLSV, WAK, and RLCK-VIIa, were previously shown to be enriched in genes with a role in biotic stress responses based on either their function or expression profile (9, 47). Thus, to investigate in greater details if xisRNA *cis*- genes coding for kinase domain proteins are likely to participate in rice responses to biotic interactions, we constructed a phylogenetic tree of *Arabidopsis thaliana* and rice kinase domain-containing proteins (Supplementary File S1). In this tree, we identified those instances where a xisRNA duplex locus was fully included within the mature transcript of a rice protein. As illustrated in Figure 4, manual examination of individual subclades of this tree with associated xisRNAs revealed some interesting insight: first, 17 xisRNA have a cis-gene product in a subtree of RLCK-VIIa proteins (Figure 4A). RLCKs physically associate with RLK receptor complexes at the plasma membrane and mediate downstream phosphorylation signaling events (11). Most of the OsRLCK clustering with RIPK, an *Arabidposis* RLCK involved in ETI signaling, stomatal defense and root development (13), have a cognate xisRNA. Still on Figure 4A, in the BIK1/PBL1 (13) clade, OsRLCK107, OsRLCK118 and OsRLCK176 have been previously shown to contribute to PTI or resistance to *Xoo* downstream of the OsCERK1, SDS2 or Xa21 receptors (48–50) and their transcript has a xisRNA duplex. Second, Figure 4B indicates that RLCK-VIIa proteins encoded by the loci overlapping with xisRNA007 (OsRLCK28) and xisRNA044 (OsRLCK55) closely cluster with the Arabidopsis decoy PBS1 (51). A *Xoo* type III effector interact with OsRLCK55 and OsRLCK185 (also in this clade) and OsRLCK185 acts as an essential phosphorelay of its binding partner OsCERK1 in mediating chitin- and peptidoglycan-induced rice immunity signaling (52). Finally, some of the most highly and broadly induced xisRNAs are associated with subgroups of RLK/Pelle protein kinase of the DLSV subfamily (Figure 4C): the xisRNA010-associated protein kinase domain is closely related to the *Arabidopsis* CYSTEINE-RICH RECEPTOR-LIKE PROTEIN KINASE 2 (CRK2) which is required for PTI and defense against a bacterial pathogen (53). xisRNA004-associated protein kinase domain is closely related to *Arabidopsis* AT1G07650 / LMK1 which possesses cell death induction activity (54). Interestingly, in a sister clade, xiRNA001-3, 20, 27 and 118 *cis*-genes correspond to rice RLCKs phylogenetically closely related to the *Arabidopsis* RLCK AT1G16670.1 / COLD-RESPONSIVE PROTEIN KINASE 1 (55).

**Figure 4:**
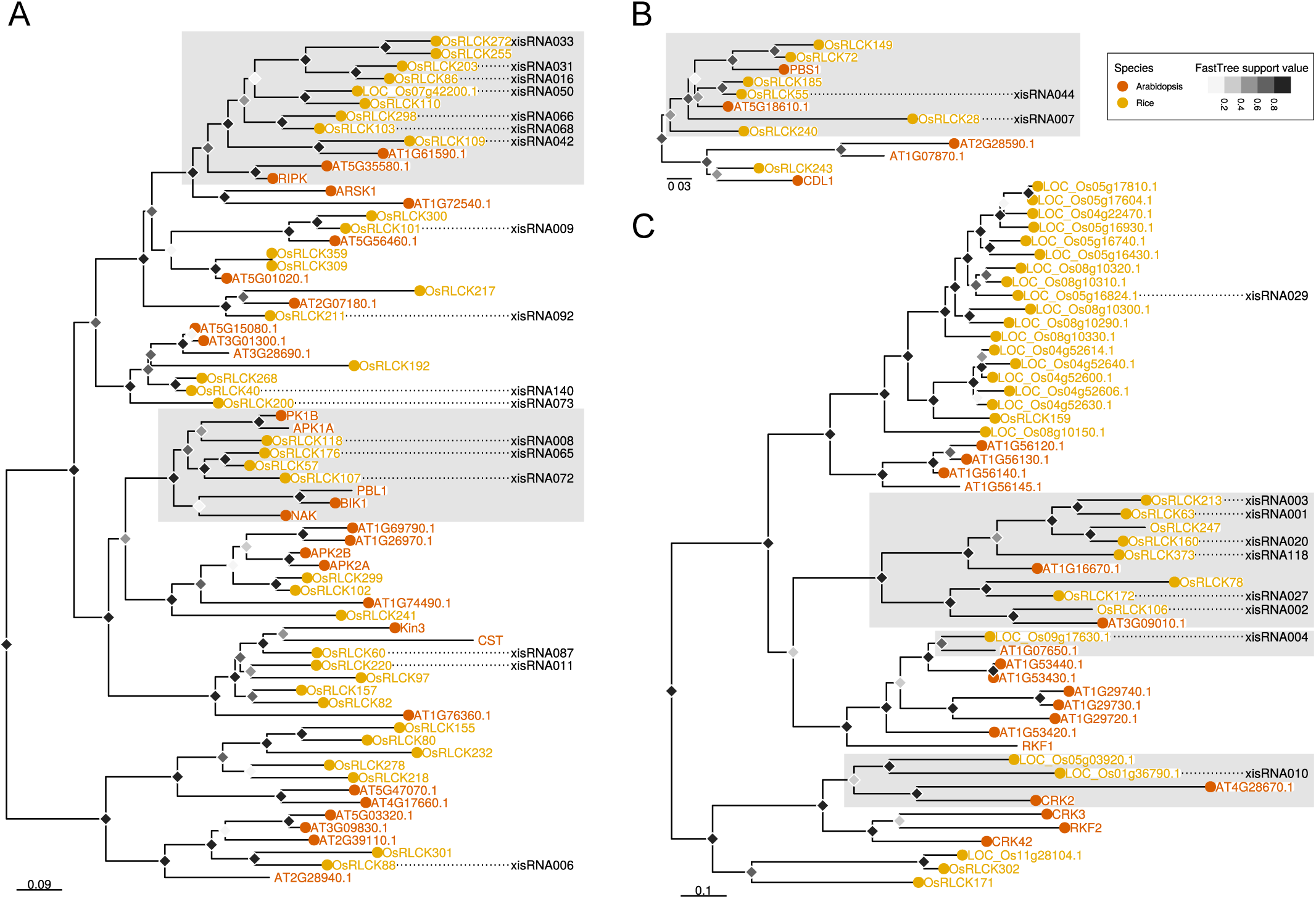
Multiple xisRNA loci cis-gene products with kinase domains are related to immune signaling pathways components in rice or Arabidopsis. The displayed phylogenetic trees correspond to selected sub-clades of proteins belonging to the RLCK-VIIa-2 (**A**), RLCK-VIIa-1 (**B**) and DLSV (**C**) subfamilies in the genome-wide phylogenetic tree (Supplementary File S1) constructed using the kinase domain of annotated proteins in the *Arabidopsis* Col-0 and Rice Nipponbare genomes. Dotted lines connect plant protein names with xisRNA identifiers when the xisRNA duplex loci is fully included in an exon of the underlying gene. Shaded rectangles highlight clades that are discussed in the text. The FastTree support value grey scale for tree nodes corresponds to local bootstrappings used by FastTree to estimate the tree’s reliability. Scale bars correspond to substitutions per site.

Overall, most xisRNA duplex loci overlap with an annotated exon. The corresponding genes are also remarkably biased for loci encoding RLK and RLCK proteins. Some protein sequences harboring a kinase domain and associated with xisRNA loci are phylogenetically or functionally linked to immune signaling. Thus, if xisRNA have regulatory activity, it is conceivable that they act in *cis* to dampen PTI signaling.

### xisRNAs accumulation is dependent on canonical components of the rice regulatory sRNAs biogenesis machinery

As a first step in characterizing the molecular mechanisms underpinning xisRNA induction, we investigated the capacity of rice lines compromised for the activity of canonical regulatory sRNA synthesis pathway components to produce xisRNAs in response to bacterial infection.

In rice, the *OsDCL3a* and *OsDCL3b* paralogs are respectively required for the biogenesis of 24nt non-based protocol canonical long miRNAs (lmiRNAs) (56) and 24nt phased small RNAs (57). In contrast, OsDCL1 is required for rice canonical ∼21nt miRNA biogenesis (58). NGS analysis of xisRNAs indicated that most of the loci generate 20-22nt long species. We therefore tested the *OsDCL1* inverted repeat transgenic line OsDCL1-IR previously produced by Zhang *et al.* (*24*) in the Nipponbare background and shown to be specifically silenced for the *OsDCL1* gene. In northern blots, the *Xoo* MAI1 strain triggered accumulation of xisRNA01 and xisRNA06 in leaves of the wild type parental control cultivar relative to water infiltrated samples, while miRNA159 levels were comparable across treatments (Figure 5A). As expected in plants with *OsDCL1* activity defects, the MAI1-inoculated OsDCL1-IR individual plants had strongly reduced miRNA159 levels. The OsDCL1-IR plants also failed to respond to MAI1-inoculations and displayed xisRNA01 and xisRNA06 levels similar to water inoculated negative controls, which suggests that *OsDCL1* is genetically required for the induction of these xisRNAs. We note that in the OsDCL1-IR individuals, the signals for the *X. oryzae* 5S rRNAs probe, acting as a proxy for MAI1 cells leaf density, tended to be lower than in Nipponbare controls. This indicates that *OsDCL1*-silenced plants may be more resistant to MAI1 infection. However, because of the OsDCL1-IR line’s reduced fertility, we were unable to obtain seeds in sufficient quantities to conduct rigorous susceptibility tests in leaf clipping assays.

**Figure 5:**
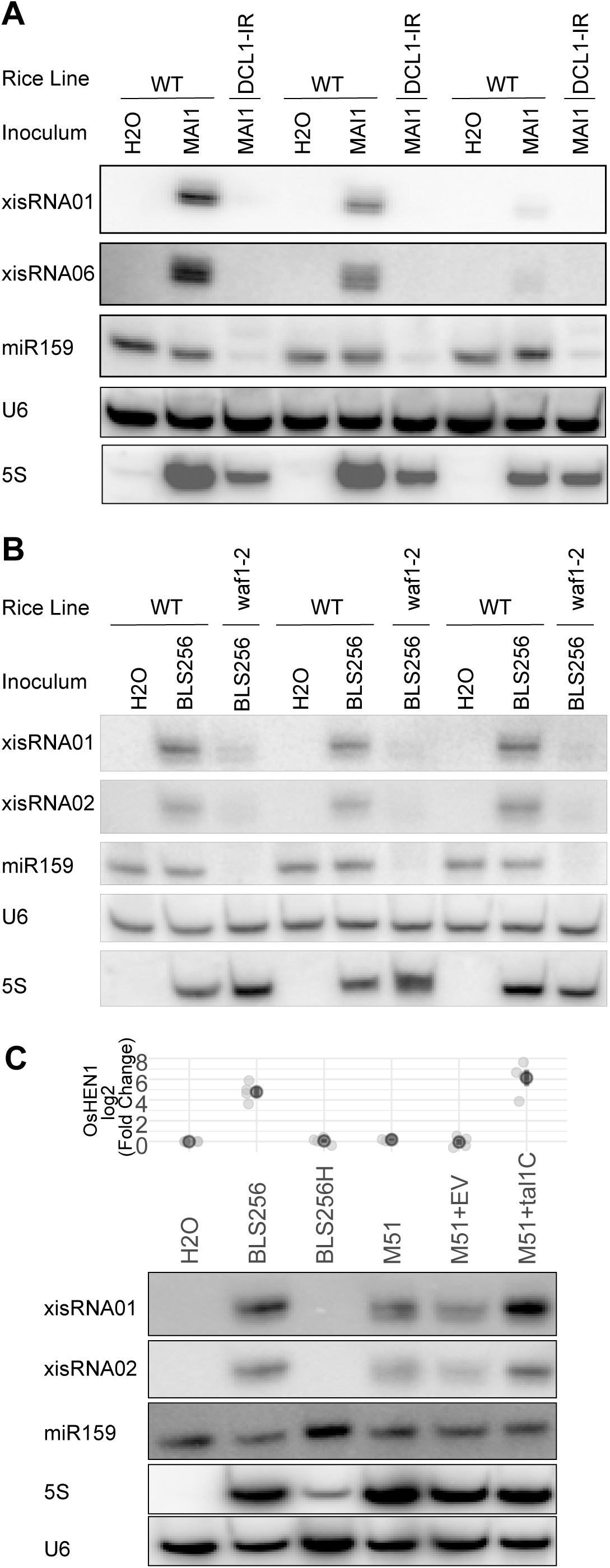
xisRNA accumulation is dependent on sRNA biogenesis genes *OsHEN1* and OsDCL1. The images of autoradiographs obtained after RNA gel northern blot analysis conducted with specific oligonucleotide probes hybridized to total RNA extracted from leaves of wild type Nipponbare (WT) or a *OsDCL1*-silenced line (DCL1-IR) in **A**, wild type Kinmaze (WT) or a *OsHEN1* mutant allele line (*waf1-2*) in **B** or Nipponbare in **C**. Leaves were infiltrated with the indicated strains and collected 24hpi. These results are representative of two and three replicate experiments for the *OsDCL1*, and the *OsHEN1* and Tal1c blots, respectively. In A and B, lanes with the same labels correspond to biological replicates of total RNA samples extracted from a single infiltrated leaf area from individual independent plants. In C, the plot on top of the gel images represent *OsHEN1* Q-RT-PCR expression data from four experiments (light gray point) with mean (dark gray points) and standard error (line ranges). The BLS256H strains correspond to a T3SS minus mutant derivative of BLS256. The “+” sign indicate that the strains were transformed with a plasmid expressing Tal1c or the empty vector (EV). For each panel, individual blot images derive from successive stripping, hybridization and detection rounds of the same membrane.

Canonical plant regulatory sRNA biogenesis pathways converge on the HEN1 methyltransferase (3). Interestingly, Both *Xoc* BLS256 and *Xoo* PXO99 strains induce the rice *OsHEN1* expression during infection as a result of the activity of a TAL effector (Tal1c and Tal9a respectively) (59). The rice *wavy leaf1-2* (*waf1-2)* line with a null mutation in *OsHEN1* has been shown to be impaired in the accumulation of miRNAs and ta-siRNA (25). This prompted us to investigate whether *OsHEN1* activity is important for xisRNAs biogenesis. To this end, we infiltrated wild type Kinmaze and *waf1- 2* individual plants with BLS256 and monitored xisRNA accumulation (Figure 5B). The miR159 probe confirmed that the *waf1-2* mutation greatly reduced miR159 accumulation in homozygous mutant individuals. While BLS256 infiltration triggered xisRNA001 and xisRNA002 accumulation in wild type plants relative to water controls, this effect was very much attenuated in the *waf1-2* individuals indicating that, in addition to *OsDCL1*, *OsHEN1* activity is necessary for xisRNA induction.

*OsHEN1* is probably universally upregulated by TALEs from Asian *X. oryzae* strains including Tal1c from the Xoc BLS256 strain and Tal9a from the Xoo PXO99 strain (59, 60). To date, the functional significance of this induction is still unknown. In separate experiments, we therefore asked whether Tal1c-mediated *OsHEN1* induction during BLS256 infection affects xisRNA accumulation. As exemplified in Figure 5C, compared to the wild type BLS256 strain, the xisRNA001 and xisRNA002 signals were slightly weaker in rice leaves infiltrated with the *tal1c* insertion mutant strain M51 (59) in the BLS256 background. Conversely, expression of *tal1c* from a plasmid in the M51 background not only restored *OsHEN1* induction but also increased xisRNA001 and xisRNA002 levels relative to M51 complemented with an empty vector. Because the effect of *tal1c* activity on xisRNA accumulation were relatively mild and to test if *tal9a* has a similar effect, we deleted the entire tal9 TALE genes cluster, which contains 5 *tal* including the *tal9a* gene, in the PXO99A strain background, as documented before (61). In an experiment summarized in Supplementary Figure S9, while this mutant had lost the ability to induce *OsHEN1* and this ability could be restored by the *Tal1c* plasmid, xisRNA023 levels were not diminished in leaves infiltrated with the tal9 cluster deleted strains relative to PXO99A. Thus, we conclude that *OsHEN1* induction during *Xoc* BSL256 infection maximizes xisRNA accumulation but a comparable effect is not observed during *Xoo* PXO99A infection.

Overall, our epistasis analysis agrees with the view that xisRNAs production requires the canonical regulatory sRNA biogenesis pathways component *OsDCL1* and *OsHEN1*. As bona fide products of these pathways, xisRNAs may thus possess PTGS-related regulatory activity.

### xisRNA accumulation is DCL4-independent but requires expression of the cis-gene

OsDCL4 is the preponderant rice DICER-LIKE responsible for the cleavage of dsRNA precursors that releases ∼21nt siRNAs associated with inverted repeat transgenes, hp-siRNAs derived from endogenous imperfect inverted repeat transcripts and ta-siRNA originating from the endogenous *TRANS-ACTING siRNA3* (*TAS3*) gene (26). As noted above and shown in Figure 6A, it was intriguing to identify the African *X. oryzae*-specific xisRNA022 duplex on the last exon of LOC_Os04g43050 which precisely corresponds to *OsDCL4*/SHO1 (8, 9). To test if *OsDCL4* is also involved in xisRNA synthesis, we examined the effect of the null *dcl4-1* allele in xisRNA accumulation in northern blot experiments. The *dcl4-1* mutation, identified in the L16S indica variety, carries a ∼1.5kb deletion (Figure 6A) and as result, does not produce *OsDCL4* transcripts (26). In these experiments, the wild type controls consisted of individuals in the progeny of selfed *dcl4-1* heterozygous parents that were genotyped as homozygous wild type at the *OsDCL4* loci whereas *dcl4-1* plants were confirmed to be homozygous for the mutant allele. The AK120922 (https://www.ncbi.nlm.nih.gov/nuccore/AK120922) transcript can form a hairpin-like structure and has been shown to be processed by OsDCL4 to produce 21nt hp-based protocol siRNAs (26). A northern blot oligonucleotide was designed to detect one of the most abundant reads on the genomic region of AK120922 in our BAI3 dataset and was used as a probe to monitor AK120922-derived hp-based protocol siRNAs. As shown in Figure 6B, while wild type and *dcl4-1* leaves expressed similar levels of miR159, the AK120922 signal detected in wild type individuals was lost in samples from the *dcl4-1* plants, thus confirming the absence of OsDCL4 activity. The *dcl4-1* mutation did not impair xisRNA01 accumulation because the corresponding signal in MAI1 infiltrated mutant plants was not decreased relative to corresponding wild type plants. In contrast, while MAI1 triggered the accumulation of xisRNA022 relative to water controls in the *OsDCL4* plants, this *OsDCL4* loci-associated xisRNA was undetectable in the MAI1-inoculated *dcl4-1* individuals.

**Figure 6:**
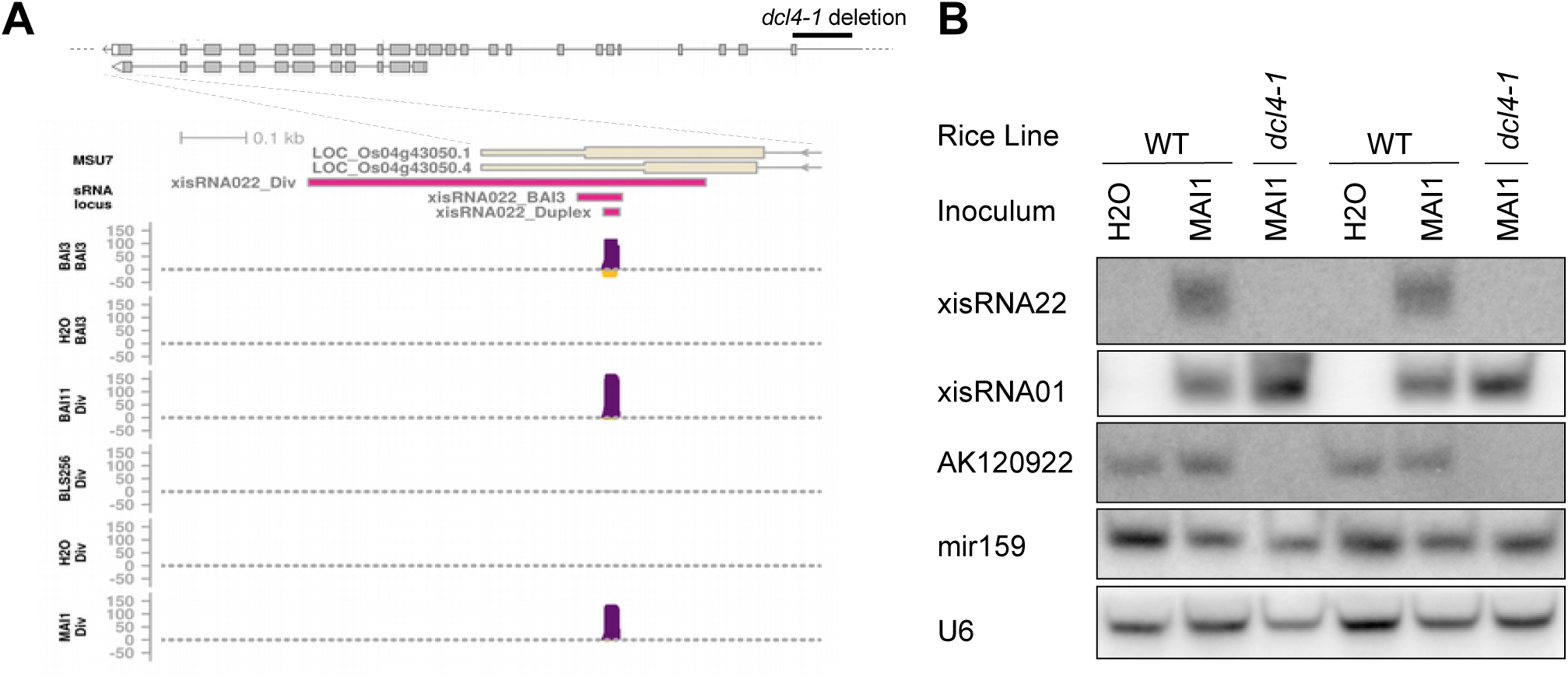
A *OsDCL4* KO allele does not impact xisRNA001 expression but compromises xisRNA022 accumulation. **A.** Diagram depicting the general exon-intron structure of *OsDCL4* and a close-up genome browser view of the last exon of the gene with the 3’ UTR. The *dcl4-1* deletion is not drawn to scale but span ∼1.5kb on the 5’-side of the gene which includes upstream promoter sequences up to the first 72bp of exon 1 (26). The genome view represents reads coverage in various treatments, the MSU7 annotation, and the xisRNA022 locus defined in the “BAI3” and “Diversity” sRNA-seq datasets as well as its duplex locus. For details see the legend of Figure 1F. **B.** Northern blot analysis conducted on total RNA extracted at 48hpi from leaves of homozygous wild type (WT) or homozygous *dcl4-1* segregants of a heterozygous parent individual. Lanes with the same labels correspond to biological replicates of total RNA samples extracted from a single infiltrated leaf area from individual independent plants. This experiment was repeated three times with similar results.

From these results, we conclude that *OsDCL4* may not play a general role in xisRNAs production. However, the *dcl4-1* null allele compromised for transcript expression is defective in the induction of its cognate xisRNA. This suggests that the expression of the protein-coding mRNA overlapping with the xisRNA duplex is likely to be required for double-stranded precursor biogenesis and consequently the production of the corresponding xisRNA.

### Genome-wide transcriptomics data provides limited support for a general role of xisRNAs in cis*-loci silencing*

We have established that xisRNA are produced at early stages of rice leaf infection relying on components of regulatory sRNA biosynthesis pathways to form duplexes of 20-22nt molecules that predominantly map to exons of protein coding genes. It is therefore conceivable that xisRNAs act in *cis* to silence overlapping transcripts. To address the potential *cis*-regulatory activity of xisRNAs, we applied a standard transcriptomics data analysis approach based on the postulate that candidate target genes of regulatory sRNA are expected to be downregulated in a treatment comparison where the corresponding sRNA is upregulated. We generated paired-end mRNA-seq data (PRJNA679478) using BAI3, BAI3H and water inoculated cultivar Nipponbare leave samples from the set of experiments that were also used to obtain the BAI3 sRNA-seq dataset. In addition, we included sequences from Nipponbare samples collected 24 hours after infiltration with the *Xoo* MAI1 strain and a water control from a previously published (62) dataset (GEO acc. GSE108504). Finally, sequences from Nipponbare samples collected 48 hours after infiltration with the *Xoc* strains BLS256, BAI11 (a.k.a. CFBP7342), or the mock control were also retrieved from another published (43) dataset (GEO acc. GSE67588). Pre-processing and downstream statistical analysis of the mRNA-seq data was performed using the ARMOR workflow (33) to compute differential expression metrics for genes annotated in the MSU7 Nipponbare annotation in comparisons involving samples inoculated with a virulent strain versus samples inoculated with the corresponding T3SS mutant strain or mock buffer (i.e. BLS256vsMOCK, BAI3vsH2O, MAI1vsMOCK, BAI11vsMOCK, BAI3vsBAI3H).

First, we examined the distribution of the log2 transform of the fold change ratio of genes in the various comparisons depending on whether an annotated transcript of the gene overlapped with a xisRNA duplex loci that is upregulated in the corresponding comparison (Figure 7A). If upregulation of a cognate xisRNA generally suppressed cis-gene expression, we would expect the median log2(FC) of this set of DE records to be inferior to zero. The computed median log2(FC) is 0.111 and, on the contrary, is significantly superior to zero (one-sided Wilcoxon rank sum test: W = 25008, p = 8e-06). Likewise, testing for a specific enrichment of genes with a cognate up-regulated xisRNA in repressed genes (log2(FC) < 0 and p-value ≤ 0.05) across treatment comparisons with a two-sided Fisher’s Exact test produced a moderately significant p-value of 0.03 for the BAI11vsMOCK comparison only. Conversely, of the 277 DE records pertaining to genes having an up-regulated cognate xisRNA in the corresponding comparison only 29 (10.5%) meet these criteria for down-regulation (Figure 7B).

**Figure 7:**
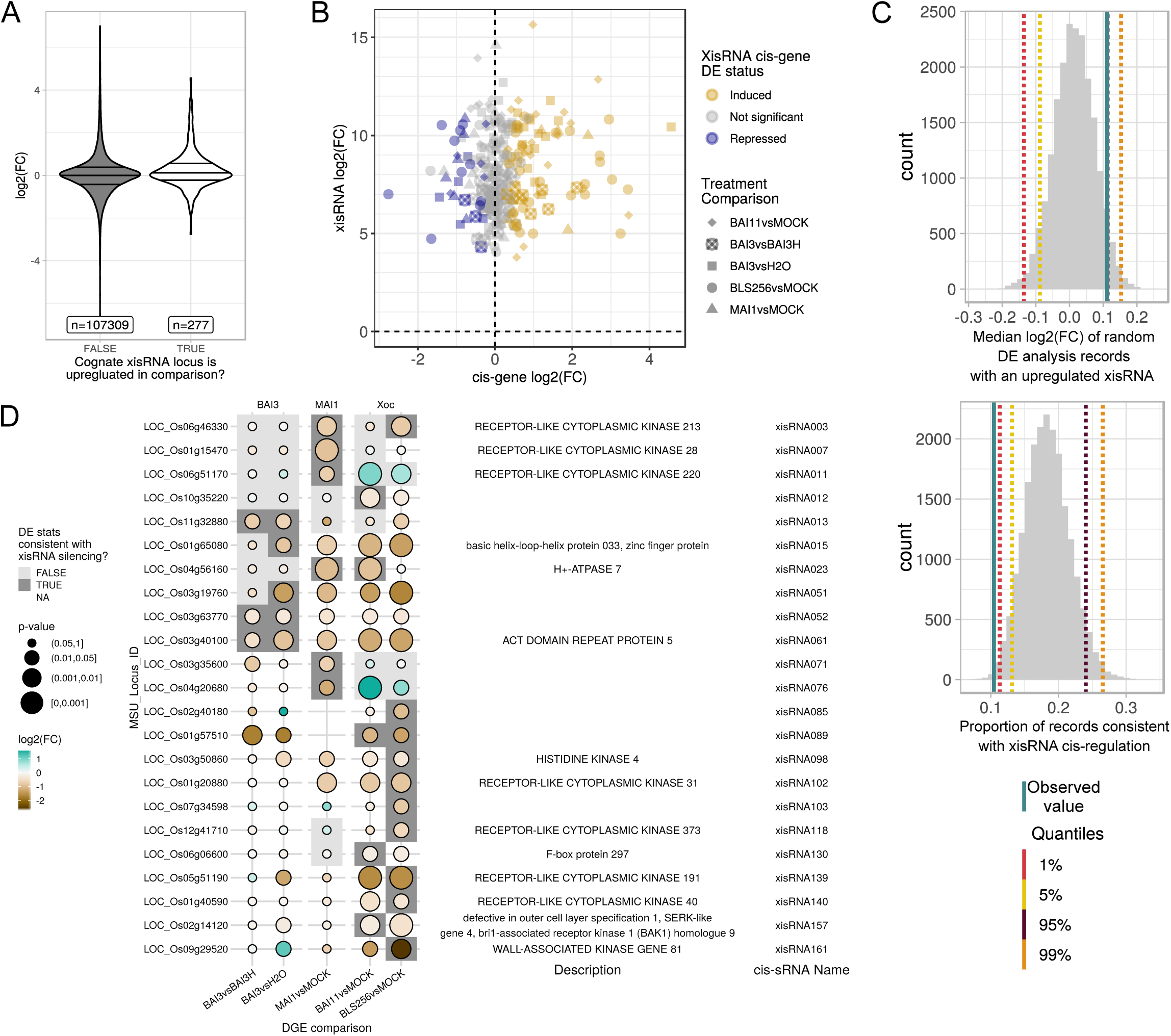
mRNA-seq data analysis points to a restricted set of xisRNA *cis*-genes as potential silencing targets. **A.** Distribution of the log_2_ transform of the fold change ratio for DE analysis records (expressed gene by treatment comparison combinations) contrasted on whether or not the gene has a cognate xisRNA and this xisRNA is also up-regulated in the same treatment comparison. The horizontal lines reflect the 25%, median and 75% quartiles. **B.** log2(FC) of xiRNA versus cognate *cis*-gene in a relevant treatment comparison. For each treatment comparison, a *cis*-gene is considered to be ‘Induced’ if it has a log2FC > 0 and a p-value ≤ 0.05, ‘Repressed’ if the log2FC < 0 and the p-value ≤ 0.05 and ‘Not significant’ otherwise. **C.** Histogram distributions of the variables computed in 20,000 trials for randomization test. **D.** Summary of the rice annotated *cis*-genes whose expression profile is consistent in at least one comparison with a possible negative regulatory effect of a cognate xisRNA. The ‘DE stats consistent with xisRNA silencing status’ is regarded as “TRUE” if in the corresponding comparison, the xisRNA has been detected as induced and if the gene DE record has log2FC < 0 and a p-value ≤ 0.05, “FALSE” if the xisRNA has been detected as induced but the gene DE record does not pass the log2FC or p-value criteria and “NA” (Not Available; white background) otherwise.

To explore further whether a xisRNA *cis*-gene gene is more likely to be down-regulated when the corresponding xisRNA is up-regulated, we performed randomization tests that consisted in 20,000 trials where each xisRNA loci was assigned a random locus identifier as cognate cis-gene. The median log2(FC) of DE records of randomly assigned genes with an upregulated xisRNA as well as the proportion of these records that are consistent with negative regulation (log2(FC) < 0 and p-value ≤ 0.05) were subsequently computed as above for the observed values. As shown in Figure 7C, 95.1% of the median log2(FC) values in the random set were lower than the observed value. Similarly, 99.4% of the proportions of records that are consistent with negative regulation in the random set were higher than the observed value.

Contrary to our initial hypothesis, this analysis of the tendency of xisRNA to suppress expression argues against the view that xisRNA generally direct *cis*-silencing. It cannot be ruled out however that xisRNA do occasionally act in *cis* to diminish transcript accumulation and we listed in Figure 7D, 23 *cis*-gene - xisRNA pairs whose DE data is consistent with a regulatory effect.

### xisRNAs duplexes exhibit limited nucleotide sequence similarity but those associated with kinase loci coincide with mRNA regions encoding a kinase active site motif

The fact that a noticeable fraction of xisRNAs is associated with genes encoding proteins with closely related kinase domains raises the question of xisRNA sequences relatedness. We performed systematic pairwise alignments between xisRNA duplex loci sequences of the most abundant read of size 19-24 mapping on the *cis-*locus antisense strand (only 148 loci considered because xisRNA133 had no antisense major reads). The resulting mismatches matrix was used to perform hierarchical clustering and derive 21 clusters of related xisRNA major reads with a count of variable positions ranging from 1 to 8 (Supplementary Figure S10). Interestingly, most individual xisRNA duplex loci in these clusters overlap a kinase encoding locus and the sequences clusters display some weak overall similarities (relatively lower mismatch counts outside the diagonal of the matrix in Supplementary Figure S10). To visualize these similarities, the multiple alignment of all reads in clusters was used to generate the consensus logo reproduced in Figure 8A. In line with our earlier observation regarding 5’ nucleotides distribution of antisense reads, major reads often start with a “TC” (or “UC” for an RNA sequence) dinucleotide on their 5’-end. The consensus pattern further points to conserved positions that seem to be phased with a 3 nt offset, reminiscent of codon code degeneracy.

**Figure 8:**
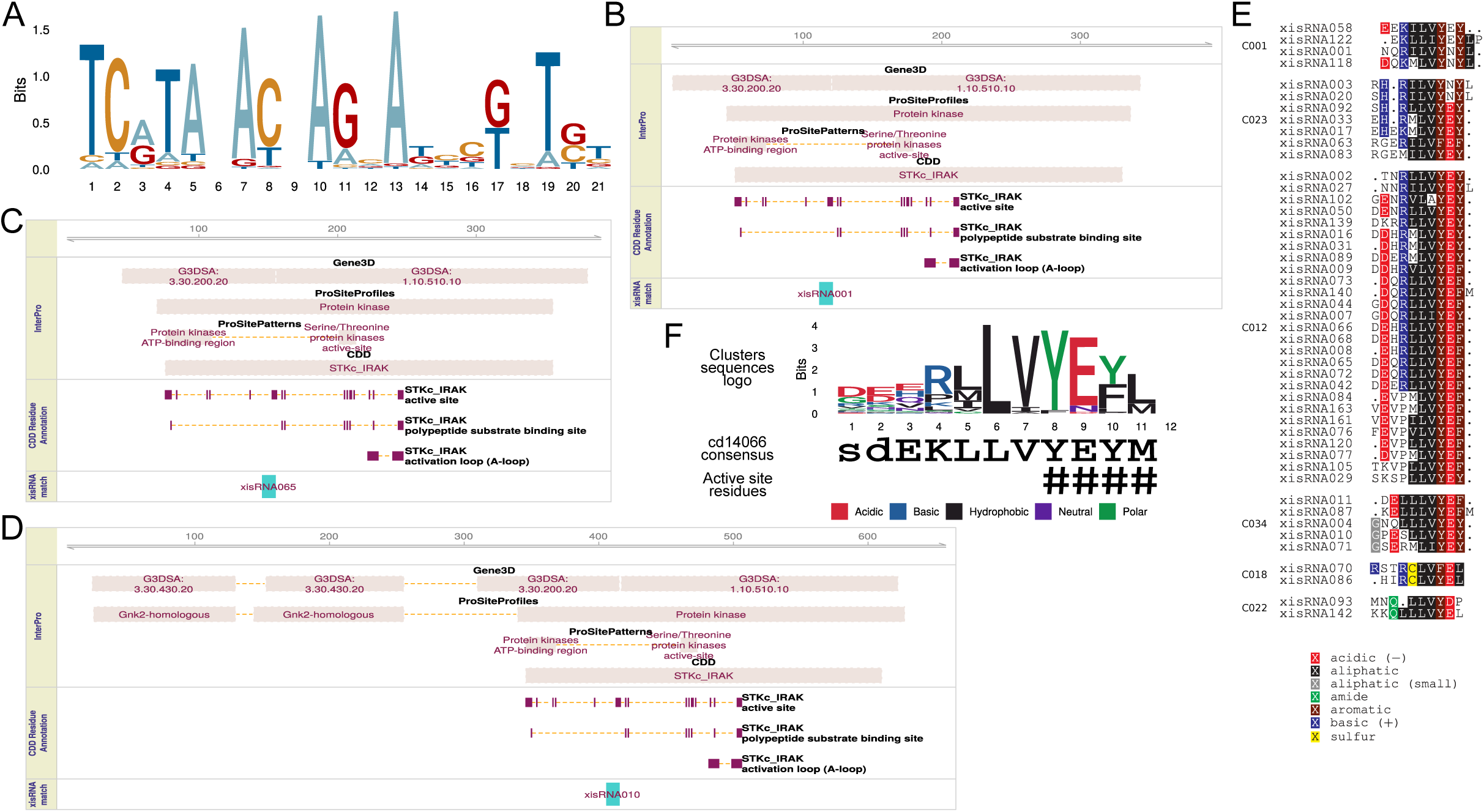
Analysis of xisRNA duplex sequences similarities. **A**. Sequence logo representation of a multiple alignment of major antisense xisRNA duplex reads sequences belonging to the clusters defined in Supplementary Figure S10. The sketches of InterPro domain hits in panel **B**, **C** and **D** for respectively xisRNA001, xisRNA065 and xisRNA010 *cis*-gene products are representative and illustrative of the observation that xisRNA duplex coding sequences consistently coincide with a specific kinase domain protein regions. **E.** Multiple protein subregion sequences alignments for a subset of 6 individual clusters from those defined and also represented in Supplementary Figure S11 (the clusters labels are consistent). **F.** Sequence logo representation of a multiple alignment of the 47 amino acid sequences displayed in E. The active site residues described for the NCBI CDD STKc_IRAK family (accession cd14066) are highlighted with hash symbols.

This pattern of conserved nucleotides in xisRNA sequences hints to the hypothesis that xisRNA duplex loci often correspond to cis-transcripts regions encoding related protein motifs. To determine if protein regions corresponding to xisRNA duplexes are likely to map to documented functional or structural domains coding sequences, we searched InterProScan hits on xisRNA duplex reference *cis*-transcripts translation products and analyzed their overlap with positions of xisRNA-associated protein subsequences (97 out of 122 that overlap an annotated transcript are also strictly within a CDS). Regardless of the provider database, most of the domains that appeared repeatedly as overlapping with a xisRNA-associated protein subsequences were connected with kinase activity. For example, 50 out of 97 overlap with a ProSite “Protein kinase” domain (PS50011). Similarly, 48 overlap with a protein structural domains CATH-Gene3D database “Phosphorylase Kinase domain 1” (<u>3.30.200.20</u>) and, as illustrated in the domain composition plots reproduced in Figure 8C-E and in the domain plots available in our dataverse, are very often positioned at a structurally conserved region of the kinase domain, aligning slightly upstream of the junction between CATH-Gene3D domains 3.30.200.20 and 1.10.510.10. As illustrated in Figure 8C-E, 39/97 xisRNA-associated protein regions overlapped with at least one residue matching with an “active site” features in the NCBI CDD database “STKc_IRAK” domain (<u>cd14066</u>). The <u>cd14066</u> CDD domain description page states that the corresponding STKc_IRAK conserved sites consensus amino acid motif ‘YEYM’ is involved in ATP binding based on structural data. As a broader complementary approach, we measured the degree of similarity between translation products of xisRNA duplex loci DNA sequences following a procedure similar to the one used above for xisRNA major reads sequences. The clustered pairwise dissimilarity scores matrix and the resulting protein sub-sequences multiple alignments are reproduced in Supplementary Figure S11. While many clusters displayed no particular similarity with one another, a subset of six clusters with a total of 47 sequences appeared to be related as judged by dissimilarity score patterns off the matrix diagonal. The vast majority of these related subsequences belongs to proteins harboring a kinase domain, many of them being classified as a RLCK (Supplementary Figure S11). In agreement with the InterPro domains overlap analysis, the consensus motif at the C-terminus of the xisRNA-associated protein sub-sequences clusters multiple alignment (LVYE[YF][LM]) match very well with the ‘YEYM’ consensus sequence of the “STKc_IRAK” ATP-binding motif in this region (Figure 8F).

In summary, our analysis of xisRNA duplex antisense sequences similarities revealed that most xisRNAs display limited sequence conservation but defined, nonetheless, a few families containing similar reads. It also appears that about half of the xisRNA duplex overlapping a CDS coincide with sequences coding for amino acid matching a conserved RLK/Pelle kinase motif that is involved in ATP binding.

## DISCUSSION

Here we report on novel 20-22nt small non-coding RNA populations highly induced by a diversity of strains of the bacterial pathogen *X. oryzae*. Only pathogenic bacteria possessing the T3SS virulence factor translocation system are fully competent for xisRNA induction.

### How well do xisRNAs fit in canonical classes of plant sRNAs?

The discovery of xisRNAs prompted an examination of rice genetic requirement for their biogenesis. xisRNA accumulation appears to utilize host regulatory small RNA synthesis pathways components *OsDCL1* and *OsHEN1* (Figure 5) but not *OsDCL4* (Figure 6). This suggests that xisRNAs derive from precursors processed by these enzymes and argues in favor of the view that xisRNAs are genuine small non coding RNAs. *OsDCL1* is dispensable for siRNA originating from CentO repeats and inverted repeats transgenes but is strictly required for the production of miRNAs (58). The ShortStack experimental sRNA loci classification algorithm did not score any of the xisRNA loci as likely *MIRNA.* Our analysis of sRNA reads properties at the xisRNA loci further indicates that upon bacterial infection most of these loci generate sRNA reads mapping to both strands of the genome sequence. These observations are incompatible with the possibility that xisRNA loci correspond to canonical *MIRNA* genes. Previous work has identified differentially accumulating rice miRNA during Asian Xoo infection (21, 22), it was therefore surprising that no xisRNA locus corresponded to a *MIRNA*. This may be due to differences in the way rice responds to Asian versus African Xoo strains or in sRNA-seq data analysis methods. Even if akin to siRNAs, xisRNA loci reads coverage supports the existence of a single xisRNA duplex at each locus (Figure 1 and 6A and our dataverse). In contrast, endogenous siRNAs loci such as hp-based protocol siRNAs, phasiRNA, transgene-derived siRNAs, or 24nt heterochromatic siRNAs generally give rise to several distinct siRNA duplexes that may be phased and that cover more or less uniformly the entire precursor sequence (5). While xisRNAs do not fit well into canonical classes of plant sRNAs, they share some features with some DCL1-dependent 21nt cis-nat-siRNAs from rice and Arabidopsis deriving from one specific site in the region of antisense transcripts overlap (63). If extrapolated to all xisRNA loci, our observation that xisRNA022 accumulation requires a sense OsDCL4-encoding mRNA (Figure 6) is consistent with this view. However, neither the reference rice Nipponbare annotation nor our BAI3-infected leaves mRNA-seq data (see coverage plots in our dataverse) argue for the widespread existence of an antisense messenger transcript complementary to the annotated protein coding mRNA. This antisense pol II RNA transcript could still be a very short-lived intermediate that does not accumulate. Alternatively, the antisense strand of the xisRNA dsRNA precursor could be the product of an RDR activity.

Our understanding of the biology of plant regulatory sRNAs bears on a few paradigmatic pathways reflecting common biosynthesis routes and regulatory activities. However, computational approaches similar to the one that have been taken here to uncover xisRNAs and relying on sRNA loci inference are revealing the hidden diversity of atypical non coding small RNA types (35, 64, 65). In this respect, rice may be a cabinet of plant sRNA curiosities because several previously unsuspected sRNA types, such as nat-miRNAs (66), 24nt phasiRNAs (67), LHR phasiRNAs (68) and lmiRNAs (56), were first described in this species.

Interestingly, the last 2-3 3’ nucleotides of xisRNA reads with a sense polarity relative to the orientation of the overlapping annotated gene show an elevated mismatch rate with the rice genome DNA sequence as compared to reads with an antisense polarity (Figure 1). This asymmetric untemplated 3’ terminal nucleotides signature seems specific to xisRNA loci (Supplementary Figure S3). The RDR2 strand of P4R2 RNAs and their DCL3-processing heterochromatic siRNA products carry a single 3’ end untemplated random nucleotide as a result of RDR2 nucleotidyl transferase activity (69). 3′ to 5′ exonucleolytic truncation and/or 3′ tailing occurs pervasively in sRNAs of an *Arabidopsis hen1* mutant but has also been detected in wild type plants (70). In this context, the tailling of unmethylated 3’ ends is accomplished by the HESO1 and URT1 uridyltransferases but other types of nuclotide additions occur in a *Arabidospsis hen1-2 heso1-2 urt1-3* triple mutant (71). The pattern of substitution in xisRNA reads untemplated 3’ nucleotides is biased in favor of cytosines (Supplementary Figure S3A). Bearing in mind that the potential caveat of biases introduced during sRNA library construction or adapter trimming cannot be formally excluded, this observation suggests that other nucleotidyltransferases that exert CC-adding activity (e.g. 72) may be involved in xisRNA tailling.

In spite of an overall predominance of 20-22nt reads, our sequencing data indicated that xisRNAs sizes vary remarkably (Figure 1C). The xisRNA bands size patterns in northern blot are consistent with sRNA-seq analysis: in addition to a major signal, minor bands were detected, reflecting the heterogeneity of xisRNAs length. DCL1 dices dsRNA precursor in homogeneously sized 21nt duplexes except for some 22nt miRNAs that trigger phasiRNA production. If xisRNAs are indeed *OsDCL1*-dependent, this raises the issue of the origin of xisRNA size variability. Just like other plant small RNAs, xisRNA require HEN1 activity for their proper accumulation (Figure 5B). The presence of noticeable proportions of reads <21nt for some xisRNA loci may indicate that they underwent 3’ to 5’ trimming. This observation and the distinctive preeminence of 3’ non-templated nucleotides in sense reads may further indicate that xisRNA undergo 3′ to 5′ exonucleolytic truncation and that 2’-O methylation by the OsHEN1 methyltransferase is a rate limiting step in xisRNA biosynthesis. The observed effect of OsHEN1 induction by Tal1c (Figure 5C) that maximises xisRNA accumulation would be consistent with this conjecture.

### Molecular activity and biological function of xisRNA

What we have learnt about xisRNA features and biogenesis argues in favor of the idea that they possess PTGS-related regulatory activity. The observed enrichment of 5’ uracyl in antisense xisRNAs (Figure 3 and Supplementary Figure S7B) is a hallmark of sRNAs loaded into OsAGO1 complexes and supports this notion. xisRNA duplex loci generally overlap with annotated rice gene bodies (91%) and are included within annotated exons of these genes (75%). However, our mRNA-seq data analysis provided little support for a general role of xisRNAs in the suppression of *cis*-gene expresssion (Figure 7). One potential pitfall of our analysis is that for the MAI1 and BLS256 strains no T3SS mutant inoculated samples were included and we thus may have overlooked PTI-regulated *cis*-genes induced by a T3SS minus strain. To demonstrate xisRNAs silencing activity, future work will need to address whether they act in *trans* through partial complementarity with target transcript and/or if their regulatory activity mostly lies in translational inhibition of target genes similar to plant 22-nt siRNAs (73).

xisRNA *cis*-genes are enriched for RLK/Pelle kinase family proteins encoding loci (Figure 3). In the absence of evidence for a *cis*- or *trans*- regulatory activity, it is difficult to interpret the significance of our results regarding the overlap between the xisRNA duplex loci in kinase genes and the coding sequence of a conserved ATP-binding site (Figure 8). miRNAs frequently target several members of gene families (2). For example, the miR482/2118 superfamily targets several plant R genes at the level of the coding sequence for a conserved P-loop region that is important for defense signalling (6). Probably evolving under different selective constraints, plant parasitic *Cuscuta* mobile sRNAs deliverd into host tissues can be grouped into superfamilies that also target conserved host protein coding sequences, thus expanding the range of accessible hosts (74). Deciding whether this enrichment of a kinase enzymatic motif coding sequences in xisRNA duplex is relevant to their biogenesis or to their activity will require an understanding of the actual nature of xisRNA molecular activity in the light of an evolutionary analysis of the diversity of these sequences in rice genomes.

We initially identified xisRNAs in experiments with the African *Xoo* strain BAI3 (Figure 1) but subsequently realized that a subset of xisRNA duplex loci are also induced by other strains of this species and that other subsets of xisRNA duplex loci seem to respond differentially depending on the genetic background of the bacterial strains (Figure 2 and 3, Supplementary Figure S5). These observations underscore the importance of this phenomenon in a disease context and implies that it is evolutionary conserved across the *Xo* species diversity and across pathovars causing two distinct diseases. Although we have not formally addressed this issue, it is likely that xisRNA are uniquely produced when *Xo* infect the rice leaf because the manual inspection of xisRNA signatures in the extensive Rice NextGen sequence database (42) found hits in no condition but in the PXO99A dataset.

The dependence on a functional T3SS for xisRNA accumulation in a compatible interaction is strong evidence that xisRNA are functionally connected to rice susceptibility but could simply be attributable to the indirect effect of the *in planta* growth defect of a T3SS strain and not to the direct and specific activity of one or several T3SS-dependent factors. However, PXO99 triggers xisRNA production in a T3SS-dependant fashion (Figure 2A) and these xisRNA are detected at a very early stage (6hpi) of leaf infection (Figure 2B). This approximately correspond to the time frame at which the earliest T3SS virulence effectors dependent effects on the host transcriptome are usually detected (75, 76). Even if some extent of *Xo* multiplication is conceivably necessary for xisRNA appearance, it is not absolutely sufficient as exemplified by several xisRNA that only respond to a subset of virulent strains (Figure 2A and Supplementary Figure S5). Consequently, we favor the hypothesis that Xo T3SS virulence effectors are directly implicated in xisRNA biogenesis.

Rice mutant lines in sRNA biogenesis pathways components suffer a range of severe reproductive and developmental phenotypes that preclude rigorous testing in classical susceptibility assays for *Xo* diseases. We showed that Tal1c-mediated *OsHEN1* induction contributes mildly to xisRNA accumulation (Figure 5C). A BLS256 derivative strain with a *tal1c* mutation is as virulent as the wildt ype (77) which can be due to an inadequacy of the assay to detect subtle phenotypes. This may also imply that this effect on xisRNA accumulation does not result in altered virulence. Our northern blot data on the *OsDCL1* silenced plants (Figure 5A) does provide some circumstantial evidence that this gene promote BLB susceptibility which is in line with previous studies on rice susceptibility to *M. oryzae* where *OsDCL1* silencing activates basal resistance to this fungus (24) while its over-expression impairs phytoalexin biosynthesis and compromises disease resistance (78).

Many of the xisRNA duplex cis-genes encoding kinase domain proteins have documented or highly likely functions in rice immune signalling (Figure 4) but, as discussed above, the level of the *cis*- transcript is rarely reduced in the presence of the cognate xisRNA. There are however exceptions, such as the xisRNA007 cis-gene that belongs to the OsRLCK185 clade (Figure 4B). Of the non-kinase loci overlapping xisRNAs, xisRNA023 is probably the most conspicuous. Not only because it is one of the most highly expressed in response to all strains except Xoc BLS256 (Figure 3), but also because its *cis*-gene, *OSA7*, which is dowregulated in the presence of xisRNA023 (Figure 7), codes for a plasma membrane H^+^-ATPase proton pomp (79) and is the rice orthologue of *Arabidopsis AHA1* and *AHA2* genes that have important functions in a range of physiological processes including immune signaling (80).

Taken altogether, we believe our results can be formalized in a working model postulating that *Xo* pathovars use T3SS-translocated virulence effectors to co-opt rice sRNA biosynthesis and activity pathways to produce an unusual class of small silencing RNAs that alter plant immune signaling in order to promote bacterial infection. This is conceptually analogous to our understanding of Cross-Kingdom RNAi in plant biotic Interaction and to the function of mobile *trans*-species sRNAs from

interacting organisms that suppress plant immunity or alter plant gene expression to favor compatibility (7). However, it is doubtful that xisRNA or their precursor are of bacterial origin because their sequences poorly match the genomic sequences of the bacterial triggers (Supplementary Figure S12). An alternative working model that appears less likely but that cannot be discounted is that rice generates xisRNAs that act in the bacterial cell to direct gene silencing and mitigate bacterial invasiveness similar to what has been reported for the bacterial plant pathogen *Pseudomonas syringae* (81) and gut bacteria (82).

This study describes the discovery of xisRNAs. Future efforts will investigate in greater details their molecular activity and biological function with the long term goal to leverage this basic knowledge to improved disease control strategies in the field.

## DATA AVAILABILITY

Deep sequencing data was downloaded from SRA using the following NCBI BioProject accessions : ‘MAI1’ mRNA-seq dataset (PRJNA427491), ‘Xoc’ mRNA-seq dataset (PRJNA280380), PXO99 infection time course sRNA-seq dataset (PRJNA252424). The PXO99A sRNA-seq dataset was downloaded from the Plant MPSS databases at https://mpss.danforthcenter.org/web/php/pages/library_info.php?SITE=rice_sRNA&showAll=true.

The sequences generated in the course of this study were deposited to SRA under the following NCBI BioProject accessions: ‘BAI3’ sRNA-seq dataset (PRJNA552731), ‘Diversity’ sRNA-seq dataset (PRJNA552775), ‘BAI3’ mRNA-seq dataset (PRJNA679478).

The R code used to generate the material described in this study as well as additional plots are stored in a dataverse and persistently available at https://doi.org/10.23708/RXIXM3.

## FUNDING

GR and AP-Q and were supported by a doctoral fellowships awarded by the French Ministry of Research and Higher Education (Ministére de l’Enseignement Supérieur et de la Recherche) and the Erasmus Mundus Action 2 PANACEA, PRECIOSA program of the European Community, respectively. JMJ was supported by an NSF Postdoctoral Fellowship in Biology (DBI – 1306196). This work was funded by the ANR with a grant (ANR-14-CE19-0009-01) to SC.

## AKNOWLEDGMENTS

The authors acknowledge the IRD itrop (https://bioinfo.ird.fr/) HPC platform at IRD Montpellier for providing HPC resources. We thank Dr. Blake Meyers and Dr. Bing Yang for granting access to their sRNA deep sequencing dataset in the Plant MPSS databases. We are particularly grateful to our colleagues who shared seeds of some of the rice genetic material used in this study: Dr. Bo Zhou for the OsDCL1-IR trangenics, Dr. Xiaofeng Cao for the *dcl4-1* line and Dr. Itoh Jun-Ichi for the Kinmaze variety and *waf1-2* mutant.

## Notes

https://doi.org/10.23708/RXIXM3

### Competing Interest Statement

The authors have declared no competing interest.

