## Supplemental_Files for "An atypical class of non-coding small RNAs produced in rice leaves upon bacterial infection": Sup_Fig_S01.pdf

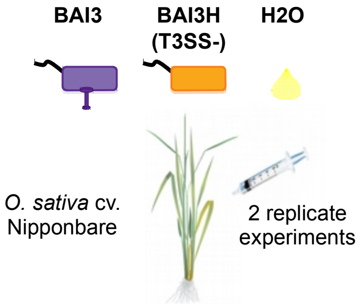


- Sample collection at 24hpi
- Total RNA extraction
- sRNA sequencing BioProject: **PRJNA552731**
- Read trimming and size selection (≥18nt)
- sRNA reads mapping: bwa aln (v.0.7.17)
- Genome segmentation (ShortStack)

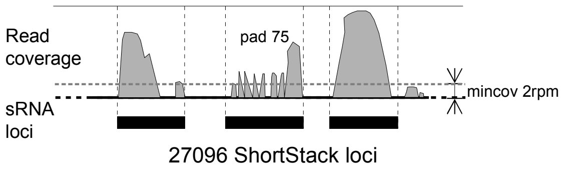


- Exclusion of tRNA and rRNA genomic loci

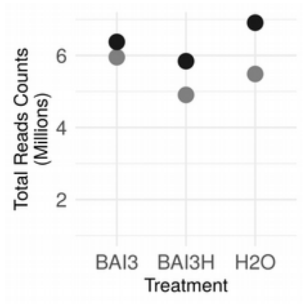


- **Differential sRNA expression (DESeq2)**
  - *- $H_0$: absolute log2 fold-change ≤ 2*
  - *- adjusted pvalue cut-off ≤ 0.05*

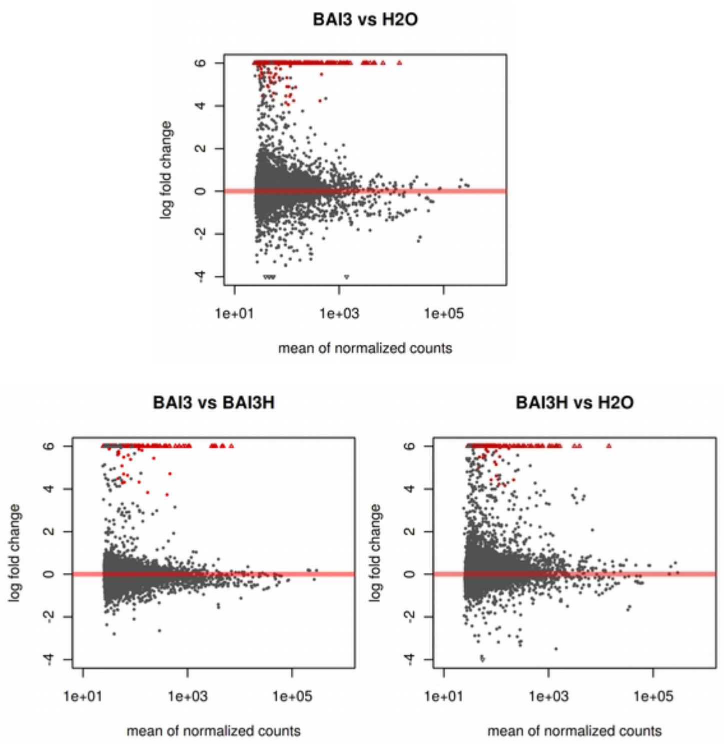


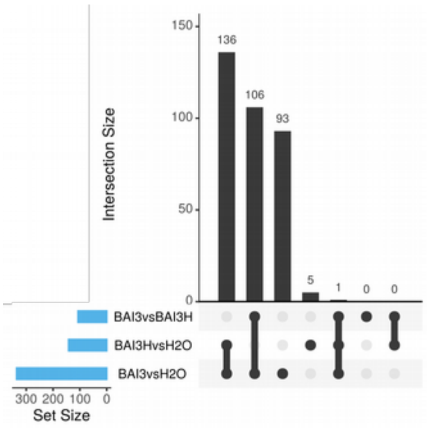


- **Xanthomonas-induced small RNAs (xisRNAs)**
  - ✓ Upregulated in BAI3vsH2O AND BAI3vsBAI3H but NOT Differentially expressed in BAI3HvsH2O
  - ✓ ≥ 75% of the reads at the locus must map uniquely to this locus
  - ✓ ≥ 50% of the reads at the locus must have length between 20 and 24nt

### 64 xisRNA loci
