## Supplemental_Files for "An atypical class of non-coding small RNAs produced in rice leaves upon bacterial infection": Sup_Fig_S06.pdf

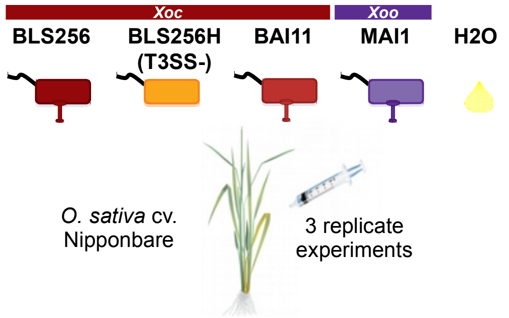


- Sample collection at 24hpi
- Total RNA extraction
- sRNA  sequencing BioProject: **PRJNA552775**
- Read trimming and size selection (≥18nt)
- sRNA reads mapping: bwa aln (v.0.7.17)
- Genome segmentation (ShortStack)

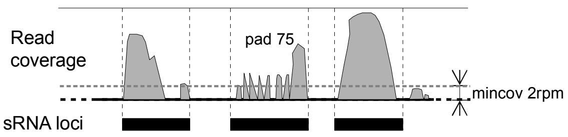


7118 ShortStack loci

- Exclusion of tRNA and rRNA genomic loci

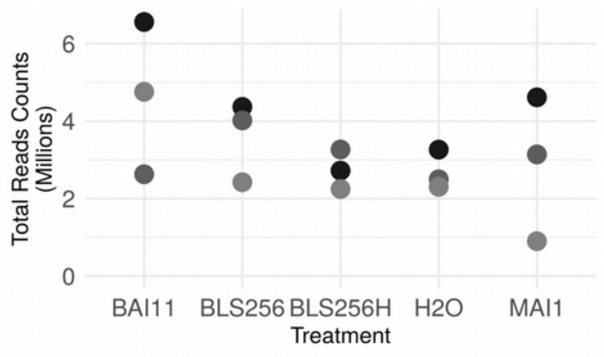


- **Differential sRNA expression (DESeq2)**
  - *$H_0$: absolute log2 fold-change ≤ 2*
  - *adjusted pvalue cut-off ≤ 0.05*

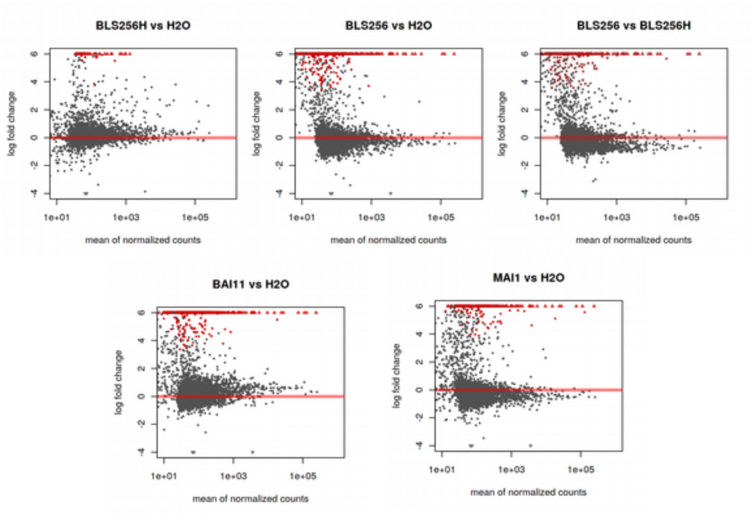


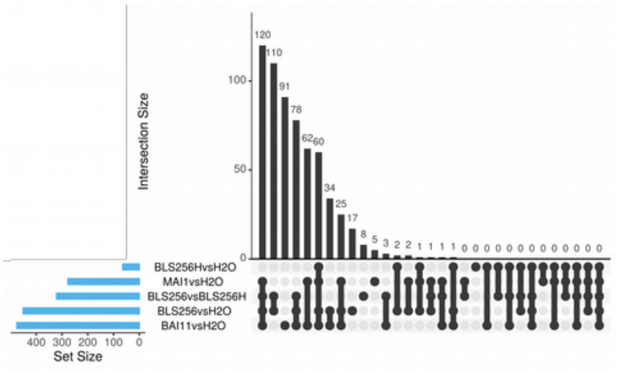


- **Xanthomonas-induced small RNAs (xisRNAs)**
  - ✓ (Upregulated in both BLS256vsBLS256H and BLS256vsH2O but NOT DE in BLS256HvsH2O) OR (upregulated in MAI1vsH2O OR BAI11vsH2O but NOT DE in BLS256HvsH2O)
  - ✓ ≥ 75% of the reads at the locus must map uniquely to this locus
  - ✓ ≥ 50% of the reads at the locus must have length between 20 and 24nt

### 141 xisRNA loci
