## Supplemental_Files for "An atypical class of non-coding small RNAs produced in rice leaves upon bacterial infection": Sup_Tables_S4-S5_and_Figures_Legends.pdf

**Supplementary Figure S1: BAI3 dataset analysis flowchart and main features**

Sketch of the overall experimental and bioinformatics procedures used for the production and analysis of the sRNA-seq data for the BAI3 dataset. Bulleted list elements correspond to the main steps in the workflow. The first dotplot depicts the total number of reads included in the DE analysis broken down by replicate sample (grey color) and treatment. The red colored symbols in the MA plots correspond to loci detected as DE in the relevant comparison. Similar to a Venn diagram, the "Upset plot" summarizes the cardinality of the intersection between various combinations of sets of DE sRNA loci in treatment comparisons.

**Supplementary Figure S2: Size distribution plots and Genome browser views for control *MIRNA* loci.**

The pri-miRNAs loci considered in this analysis correspond to those reference rice sRNA loci annotations in the Plant small RNA gene annotations database (1) that are flagged as "MIRNA" and for which an overlapping experimental sRNA loci has been identified in all the datasets considered in this study. The Dicer Call (DC) flag values (20, 21, 22 or 24) included in the database are taken as indicative of the overall general size of these miRNAs in rice libraries.

**A**. Distribution of the the fraction of reads by length for pri-miRNAs loci as a function of their Dicer Call in the Plant small RNA database. One data point is the proportion of reads of the corresponding length mapped to an individual pri-miRNAs loci and deriving from libraries constructed with BAI3 inoculated leaves.

**B**. Similar to A but represents size proportions as stacked bar plots for selected pri-miRNAs loci. The total number of reads from the libraries considered to compute the proportions at individual loci are printed on the bars. Columns in the plots grid group together pri-miRNAs loci by Dicer Call values and the rows correspond to the experimental treatment.

**C**. Genome browser snapshots of a few selected pri-miRNA loci with their miRBase annotation and read coverage tracks (reads per million) for the various experimental treatments. Coverage of reads mapping to the top genomic strand is represented with positive values and a violet colored area whereas coverage of reads maping to the opposite strand is represented with negative values and a golden colored area.

**Supplementary Figure S3: BAI3 xisRNA reads mapping to the same strand as the annotated transcript exhibit specific 3'-end mismatches.**

**A**. Nucleotide substitution matrix of mismatches for positions -1 and -2 relative to the 3'-end of reads from the BAI3 treatment mapping to a xisRNA loci and whose polarity is the same as the annotated *cis*-gene. The counts correspond to the total number of read *vs* genome nucleotides comparisons that yielded the corresponding substitution pattern. Comparisons where nucleotides matched are not included in the matrix.

**B**. Distribution of genomic sequence mismatch ratios along relative positions within reads for control experimental sRNA loci. This plot is similar to panel I of Figure 1 although in this case the mismatch ratios were computed for a set of 190 control experimental sRNA loci defined in the BAI3 dataset. They were selected based on the following criteria: they are not xisRNA loci, they overlap with a rice sRNA loci of the Plant small RNA gene annotations database (1), they overlap on at least 20bp with an MSU7 annotated exon, their ShortStack Dicer Call is between 20 and 23, and their ShortStack 'FracTop' values is between 0.1 and 0.9. Each data point represents the ratio of the number of mismatches over the total number of comparisons with the genomic sequence for that position of all the reads from the BAI3 treatment and mapping to a sRNA loci of the control set in the sense orientation (violet) or antisense orientation (golden) relative to the MSU annotated gene overlapping with the sRNA loci. None of the two-sided Wilcoxon signed rank tests measuring differences in sRNA mismatch ratios between read orientations performed at each position were significant (BY adjusted p-value ≤ 0.05). The bar graph on the top reports on the total count of comparisons for each orientation.

**C**. Similar to panel B except that the colored points represent mismatch ratios for the same control *MIRNA* loci as those described for Supplementary Figure 2A (golden) versus xisRNA loci defined in the BAI3 data set (pink). Here, only reads from the BAI3 treatment mapping to the sense strand of the *cis*-gene or the MIRNA loci are considered. Positions where a one-sided Wilcoxon signed rank tests measuring if the mismatch ratios for xisRNA loci is greater than for control MIRNA loci was significant (BY adjusted p-value ≤ 0.05) are marked with a "*" sign. Here, "***" indicate an adj. P-val. ≤ 0.001.

**Supplementary Figure S4: Northern blots of a xisRNA accumulation time course experiment.**

Autoradiographs obtained after RNA gel northern blot analysis conducted with specific oligonucleotide probes hybridized to total RNA extracted from Nipponbare leaves infiltrated with the indicated strains and collected at the indicated time points (hpi = hours post inoculation). The BAI3H strains corresponds to a T3SS defective mutant derivative of the BAI3 virulent strain. The xisRNA probes detect RNAs complementary to the sense strand of the xisRNA *cis*-gene. The two columns of images were obtained from two membranes having samples collected from the same infiltration experiment and ran on two separate gels for migration and transfer. Blotting, hybridization and signal detection were performed at the same time and in the same conditions. For a given probe, images from both blots are displayed with the same rendering settings (contrast and intensity). Individual images in each column derive from successive stripping, hybridization and detection rounds of the same membrane. This experiment was performed once.

**Supplementary Figure S5: xisRNAs are induced by diverse *Xoo* and *Xoc* strains.**

Autoradiographs obtained after RNA gel northern blot analysis conducted with specific oligonucleotide probes hybridized to total RNA extracted from Nipponbare leaves infiltrated with the indicated strains and collected 48h post inoculation. The BAI3H strains corresponds to a T3SS minus mutant derivative of the BAI3 virulent strain. The xisRNA probes detect RNAs complementary to the sense strand of the xisRNA *cis*-gene. The two columns of images were obtained from two membranes having samples collected from the same infiltration experiment and ran on two separate gels for migration and transfer. Blotting, hybridization and signal detection were performed at the same time and in the same conditions. For a given probe, images from both blots are displayed with the same rendering settings (contrast and intensity). Individual images in each column derive from successive stripping, hybridization and detection rounds of the same membrane. This experiment was performed once.

**Supplementary Figure S6: Diversity dataset analysis flowchart and main features.**
Sketch of the overall experimental and bioinformatics procedures used for the production and analysis of the sRNA-seq data for the Diversity dataset. Bulleted list elements correspond to the main steps in the workflow. The first dotplot depicts the total number of reads included in the DE analysis broken down by replicate sample (grey color) and treatment. The red colored symbols in the MA plots correspond to loci detected as DE in the relevant comparison. Similar to a Venn diagram, the “Upset plot” summarizes the cardinality of the intersection of various combinations of sets of DE sRNA loci in treatment comparisons.

**Supplementary Figure S7: Joint analysis of the BAI3 and Diversity sRNA-seq datasets**.
**A.** Upset plot of the cardinality of the intersection of various combinations of sets of DE xiRNA loci in treatment comparisons across the BAI3 and diversity datasets.
**B.** Distribution of the proportion of 5’ nucleotides at the first position of the reads from the BAI3 and the Diversity datasets computed for individual xisRNA duplex loci depending on whether the reads map to the sense or the antisense strand of the *cis*-gene. Note that for xisRNA loci that do not overlap with an annotated gene, reads mapping to the top strand were arbitrarily considered to be in the sense orientation. The horizontal dashed lines indicate the background proportion of the corresponding nucleotide calculated using the whole rice Nipponbare nuclear genome.

**Supplementary Figure S8: Gene Ontology terms enrichment in xisRNA-associated genes.**
Only those GO Slim terms with an enrichment test adjusted p-value ≤ 0.05 are displayed.

**Supplementary Figure S9: The tal9 region of PXO99A does not contribute to xisRNA accumulation.**
*OsHEN1* induction Q-RT-PCR data (bar plot) and sRNA accumulation (northern blots) produced with total RNA extracted from Nipponbare leaves infiltrated with the indicated strains and collected

24h after inoculation. The Δtal9 strains correspond to two independent colonies produced by deleting the tal9 cluster in PXO99A. The “+” sign indicate that the strains were transformed with a plasmid expressing *Tal1c* or the empty vector (EV). This experiment was performed once.

**Supplementary Figure S10: Sequence similarities among xisRNA major reads**

**A**. Matrix of distances as computed by the nmismatch function of the Biostrings R package for pairwise global alignments between xisRNA major reads. A ‘major read’ sequence is the most abundant unique sequence of size between 19 and 24nt from those mapping to a xisRNA duplex loci (at least 15nt included in the duplex) overlapping an annotated MSU7 gene and corresponding to the antisense strand of this *cis*-gene. Rows and columns are ordered based on a Hierarchical cluster analysis. xisRNA sequence clusters membership was determined by cutting the hierarchical tree in order to generate 115 clusters. Clusters that contained at least two xisRNA major reads were labeled (eg C025) on the left and on top of the heatmap. The “Kin. Fam.” annotation column reflects the kinase subfamily of the xisRNA locus *cis*-gene.

**B**. Multiple sequence alignments of the xisRNA major reads within the clusters defined in A.

**Supplementary Figure S11: Amino acid sequence similarities of the subregions encoded by xisRNA duplex overlapping coding sequences.**

xisRNA duplexes genomic intervals were transposed into positions in the coding sequence space of overlapping annotated transcripts to obtain approximate intervals in protein sequence by mapping CDS positions to codon number. As in Supplementary Figure 9, a matrix of distances and a Hierarchical cluster analysis were performed using the amino acid sequences of the corresponding protein subregions. Pairwise global alignment distances were computed with the dis function of the bios2mds R package. xisRNA-associated protein subsequences cluster membership was determined by cutting the hierarchical tree in order to generate 44 clusters. Clusters that contained at least two xisRNA-associated subsequences were labeled (eg C001) on the left and on top of the heatmap. Multiple sequence alignments of the amino acid sequence within these clusters is displayed on the left of the heatmap. The “Kin. Fam.” annotation column reflects the kinase subfamily of the xisRNA locus *cis*-gene.

**Supplementary Figure S12: The sequence identity of xisRNA major reads with their blast hits in *Xo* genomes sequences.**

The blastn executable was used to search hits for all xisRNA major reads sequences (see definition in the legend of Supplementary Figure 9) in the *Xo* genomes listed on the x-axis. The proportion of identical nucleotide (y-axis) was computed by dividing the values of the blast results field "nident” by those of the “qlen” field and only the highest value per query sequence and subject genome was retained to compute the distributions displayed in the box plot.

**Supplementary Table S1: List of the differentially expressed experimental sRNAs loci in the BAI3 dataset.**
A short description of the content of the columns is provided as follows:
seqnames: rice chromosome name
start: starting position on chromosome
end: end position on chromosome
width: span of the loci in bp
strand: sRNA locus strand as defined by ShortStack
gbrowse_location: genomic location of the loci formated for easy copy and paste in a genome browser.
Name: label of the loci
For the “FracTop”, “MajorRNA”, “MajorRNAReads”, “Complexity”, “DicerCall”, “MIRNA”and “PhaseScore” columns, please refer to the ShortStack manual for a description of these values.
OlapGene_Names: MSU identifier of sRNA loci overlapping annotation elements
OlapGene_Note: MSU description of sRNA loci overlapping annotation elements
OlapSimpleTrx_Names: Identifier of the MSU locus if the xiRNA locus overlap with an annotated transcript.
DEpattern_BAI3vsH2O_BAI3vsBAI3H_BAI3HvsH2O: binary string encoding the differential expression call in the comparisons listed in the column name; 1 indicates that the sRNA locus was designated as differentially expressed.
%01_Hit.s.: percentage of reads that map to a single best location on the rice genome
%02_Hit.s.: percentage of reads that map to two locations with maximal best scores on the rice genome
%20_nt: percentage of reads of size 20nt
%21_nt: percentage of reads of size 21nt
%22_nt: percentage of reads of size 22nt
%23_nt: percentage of reads of size 23nt
%24_nt: percentage of reads of size 24nt
EditDistance_00: percentage of reads with an edit distance equal to 0 relative to the genome sequence
EditDistance_01: percentage of reads with an edit distance equal to 1 relative to the genome sequence
EditDistance_02: percentage of reads with an edit distance equal to 2 relative to the genome sequence
pri_miRNA: name of overlapping pri-miRNA locus
xisRNAName: name of the corresponding xisRNA
rpm_BAI3: average expression in rpm in the BAI3 treatment
rpm_BAI3H: average expression in rpm in the BAI3H treatment
rpm_H2O: average expression in rpm in the H2O treatment
Column names with a “de_” prefix correspond to the DESeq2 output. Please refer to the DESeq2 manual for a description of these values.

**Supplementary Table S2: List of the differentially expressed experimental sRNAs loci in the Diversity dataset.**
See the legend of Supplementary Table S1

**Supplementary Table S3: xisRNAs features.**
Column names are self-explanatory and refers to the values depicted graphically in Figure 3.

### Supplementary Table S4: *X. oryzae* strains used in this study.

| Strain Designation | Other Designation | Description | Source |
|---|---|---|---|
| BAI3 | BAI3$^R$ | A rifampicin resistant and fully virulent derivative of the BAI3 wild type *Xoo* strain from Burkina Faso | (9) |
| BAI3H | BAI3$^R$ΔhrcC | A T3SS minus mutant derivative of the BAI3 strain | (9) |
| BAI15 | | Wild type *Xoc* isolate from Burkina Faso (Bagré) | (10) |
| BAI20 | | Wild type *Xoc* isolate from Burkina Faso (Cascades) | (10) |
| BLS256 | | Wild type Philippine Xoc isolate | (4) |
| BLS256H | | A T3SS minus mutant derivative of BLS256 | (5) |
| GXO1 | | Wild type *Xoc* isolate from China (Guangxi) | (11) |
| GXO7 | | Wild type *Xoc* isolate from China (Guangxi) | (11) |
| KACC10331 | | Wild type *Xoo* isolate from Korea | (12) |
| M51 | | Single marker exchange mutant in the *tal1c* gene of BLS256 | (6) |
| M51 + tal1c | | M51 harboring the pKEB31-*tal1c* (pCS472) plasmid which is a pKEB31 containing the *tal1c* gene of *Xoc* BLS256 | (7) for the plasmid and this study for the strain. |
| M51 + EV | | M51 harboring the pKEB31-*talΔCRR* (pAC99) plasmid which is a pKEB31 containing the *tal1c* gene of *Xoc* BLS256 deprived of the central repeat domain coding sequence | (7) for the plasmid and this study for the strain. |
| MAI1 | | Wild type Xoo isolate from Mali (Niono) | (8) |
| MAI10 | | Wild type *Xoc* isolate from Mali (Niono) | (8) |
| MAI18 | | Wild type *Xoc* isolate from Mali (Office du Niger) | (10) |
| MAI20 | | Wild type *Xoc* isolate from Mali (Office du Niger) | (10) |
| PXO99A | | 5-Azacytidine resistant derivative of the race 6 Filipino strain PXO99 | (2) |
| PXO99H | PXO99A ME7 | A T3SS defective mutant derivative of the PXO99A strain | (3) |
| X11-5A | | *Xo*-like strain isolated in the USA | (13) |

### Supplementary Table S5: Oligonucleotide used in this study.

| Primer sequence (5’->3’) | Target | Reference | Use |
|---|---|---|---|
| AACCAAAGAATCCTTGTCTACAATTATCTT | xisRNA001 | This study | LMW northern blots |
| TAACAGACTACTAGTGTATGAATA | xisRNA002 | This study | LMW northern blots |
| GGGGCGTACAGCGTCTCCTAG | xisRNA006 | This study | LMW northern blots |
| TCAGCGCCTGCTTATTTATGA | xisRNA007 | This study | LMW northern blots |
| AGAGAGCCTGCTTGTTTATGA | xisRNA010 | This study | LMW northern blots |
| TGACGAACTACTTCTTGTGTACGAGTTC | xisRNA011 | This study | LMW northern blots |
| AGTGATATTTGTTTGTCCAGAGATA | xisRNA022 | This study | LMW northern blots |
| CAGCAGAACTACACAGTGTGAGGAAC | xisRNA023 | This study | LMW northern blots |
| CCTGAAAGTTCATGAAGCGGT | AK120922 | This study | LMW northern blots |
| TGTATCGTTCCAATTTTATCGGATGT | U6 snRNA | This study | LMW northern blots |
| CCTGGCGATGACCTACTCTC | *X. oryzae* 5S ribosomal RNA | This study | LMW northern blots |
| CAGAGCTCCCTTCAATCCAAA | miR159 | This study | LMW northern blots |
| TCTCGGAAGGGTCTCACACGA | *OsHEN1* | This study | Q-RT-PCR |
| AGAGCACAGCCGTTGGAACAC | *OsHEN1* | This study | Q-RT-PCR |
| GAAGTCTCATCCTACCTGAAGAAG | *OsEF-1*α | (14) | Q-RT-PCR |
| GTCAAGAGCCTCAAGCAAGG | *OsEF-1*α | | Q-RT-PCR |
| CAGACGGTGTATGTAACACC | *OsDCL4* and *dcl4-1* forward | (15) | *dcl4-1* genotyping |
| CGGACTCCAAGACGCAATATGT | *OsDCL4* reverse | (15) | *dcl4-1* genotyping |
| TTGCGATCACCACAGCTTGC | *dcl4-1* reverse | (15) | *dcl4-1* genotyping |
| AAGGAGAGACTCAGAGCTTC | *OsHEN1* forward | (16) and Itoh Jun-Ichi personnal communication | *waf1-2* CAPS marker (EcoRV cleavage is diagnostic of the *waf1-2* allele) |
| ATGAGAACAGAAGGTGTGCC | *OsHEN1* reverse | | *waf1-2* CAPS marker (EcoRV cleavage is diagnostic of the *waf1-2* allele) |
