## Supplementary figures and images for "An atypical class of non-coding small RNAs produced in rice leaves upon bacterial infection"

### Sup_Fig_S02.pdf

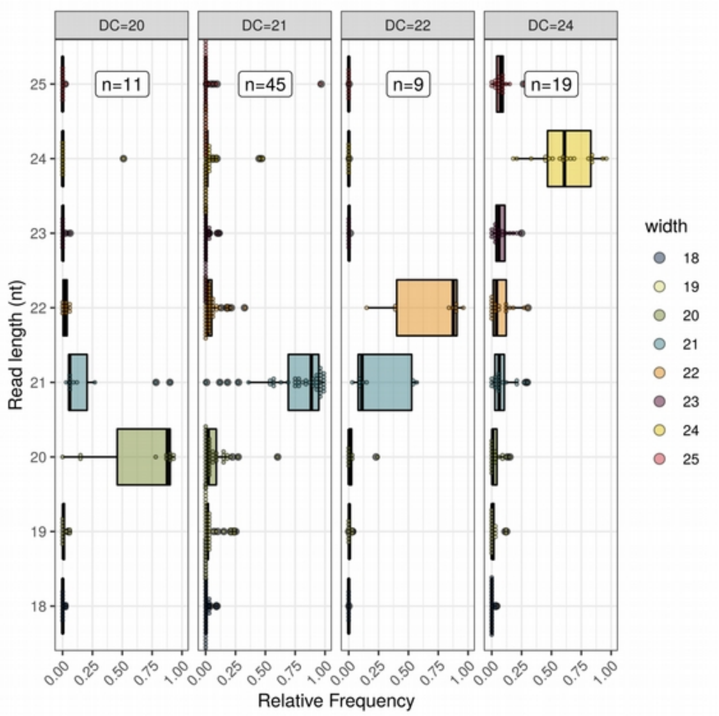

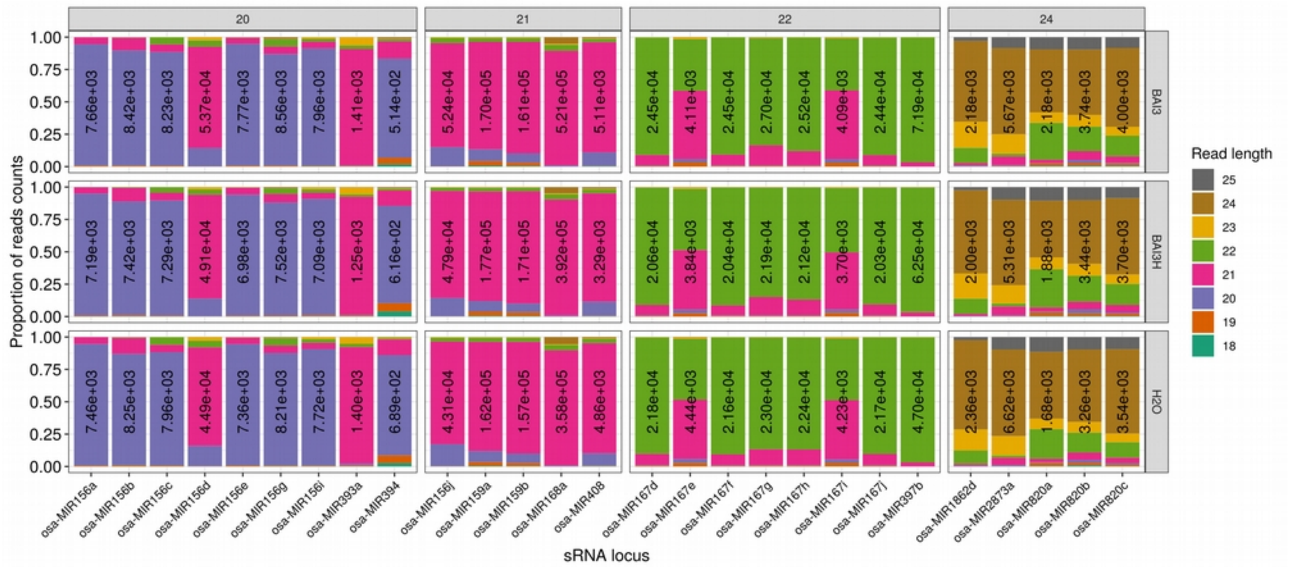

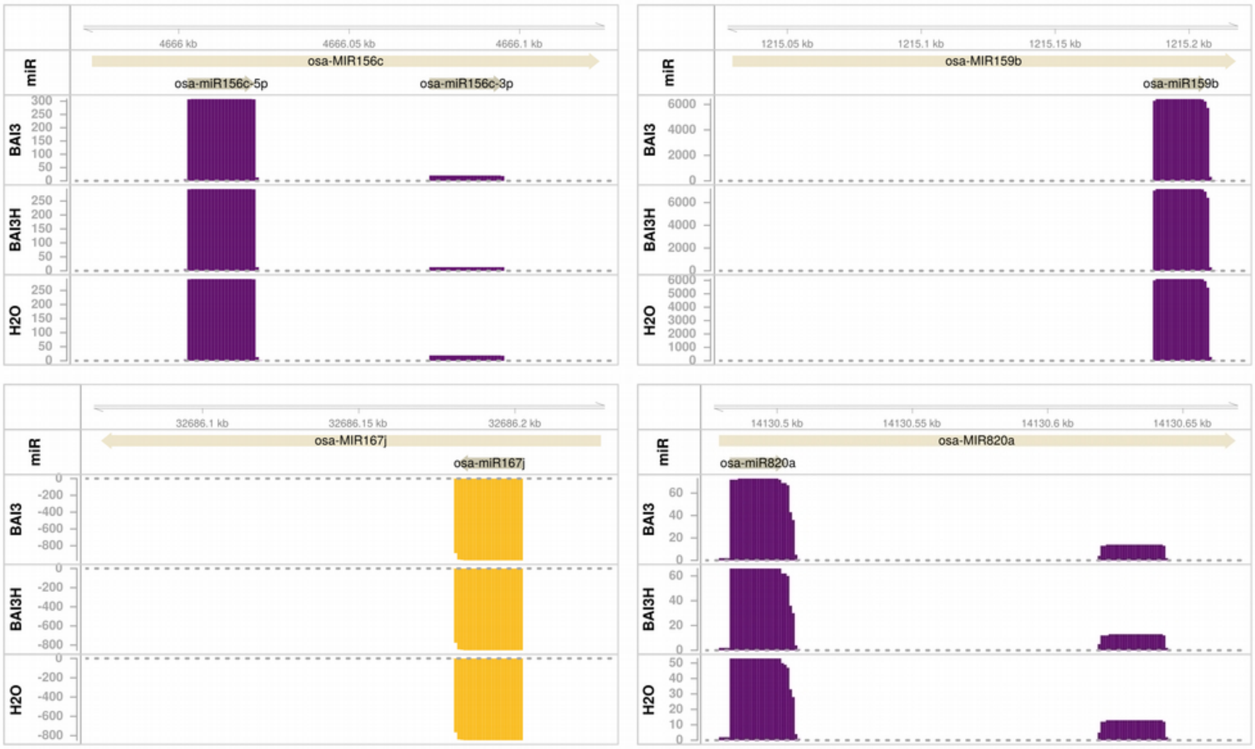

### Sup_Fig_S03.pdf

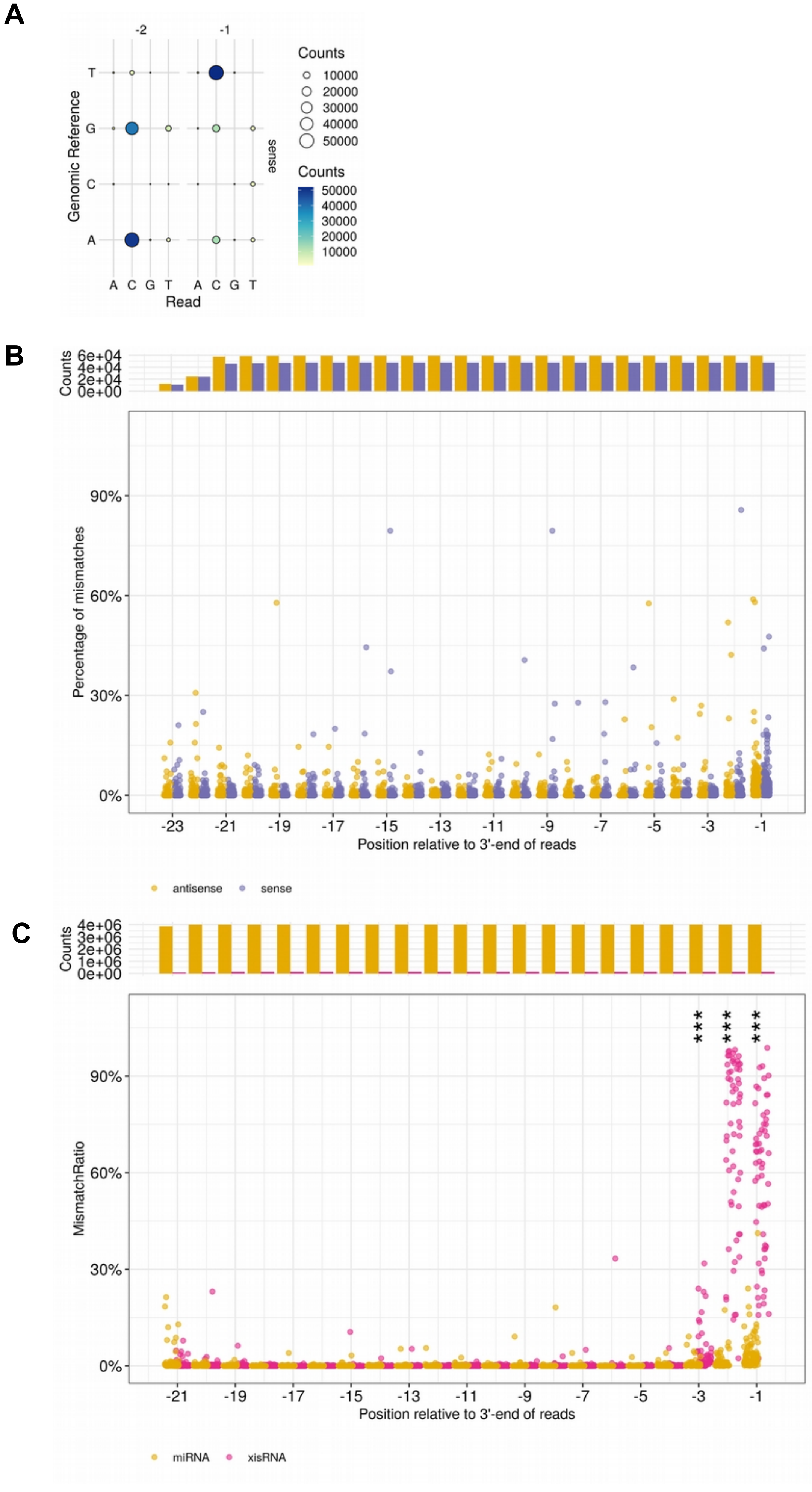

### Sup_Fig_S04.pdf

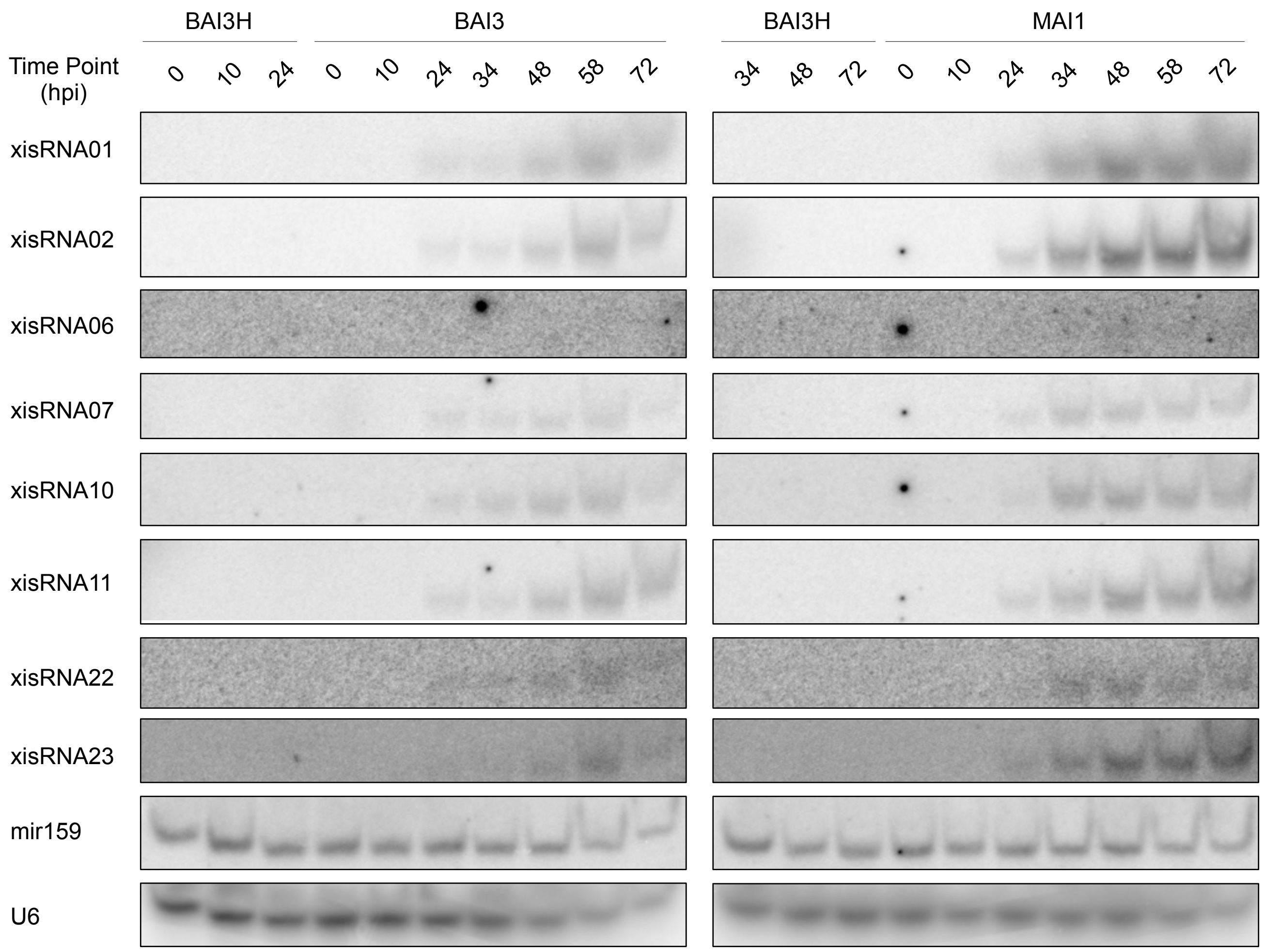

### Sup_Fig_S05.pdf

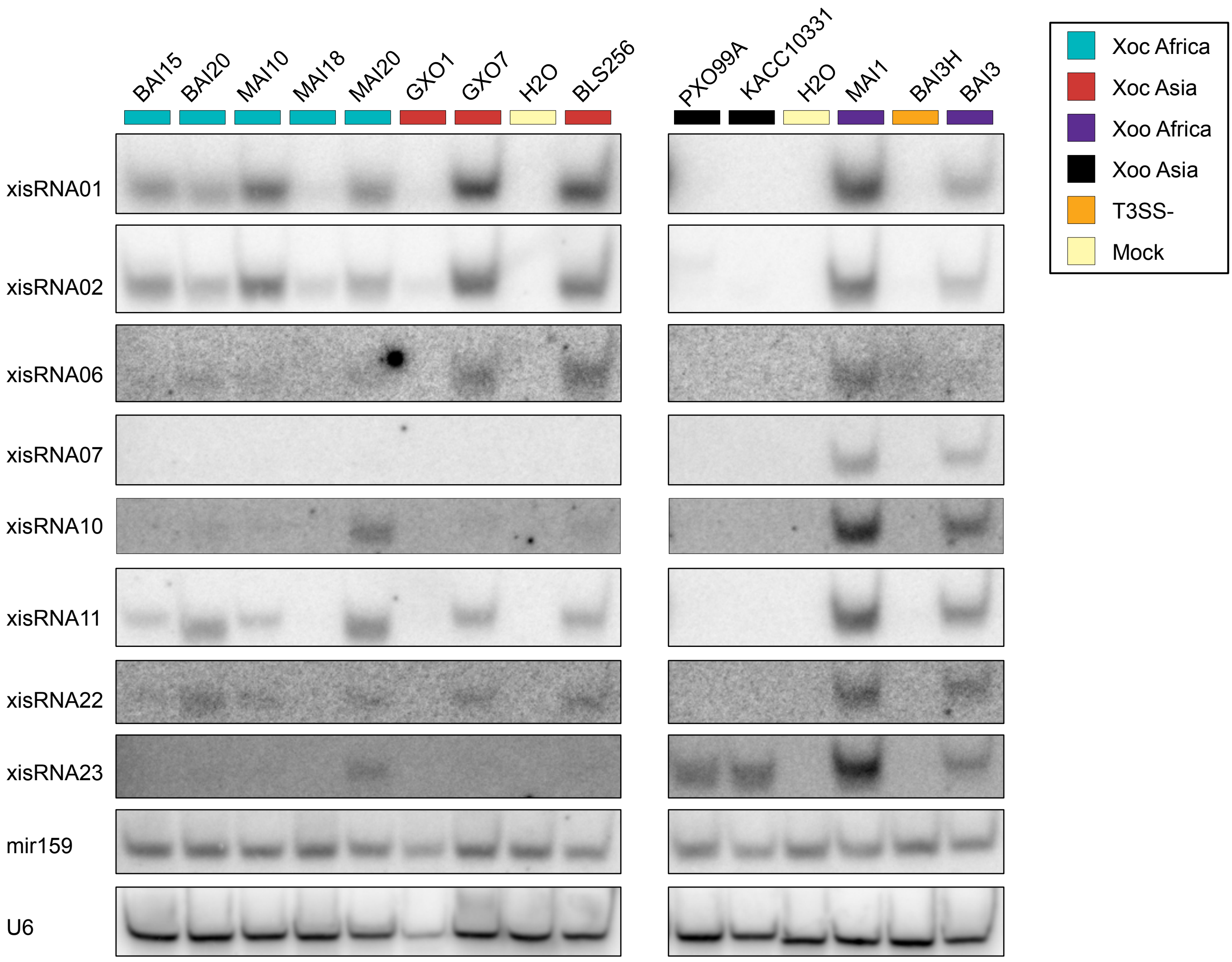

### Sup_Fig_S07.pdf

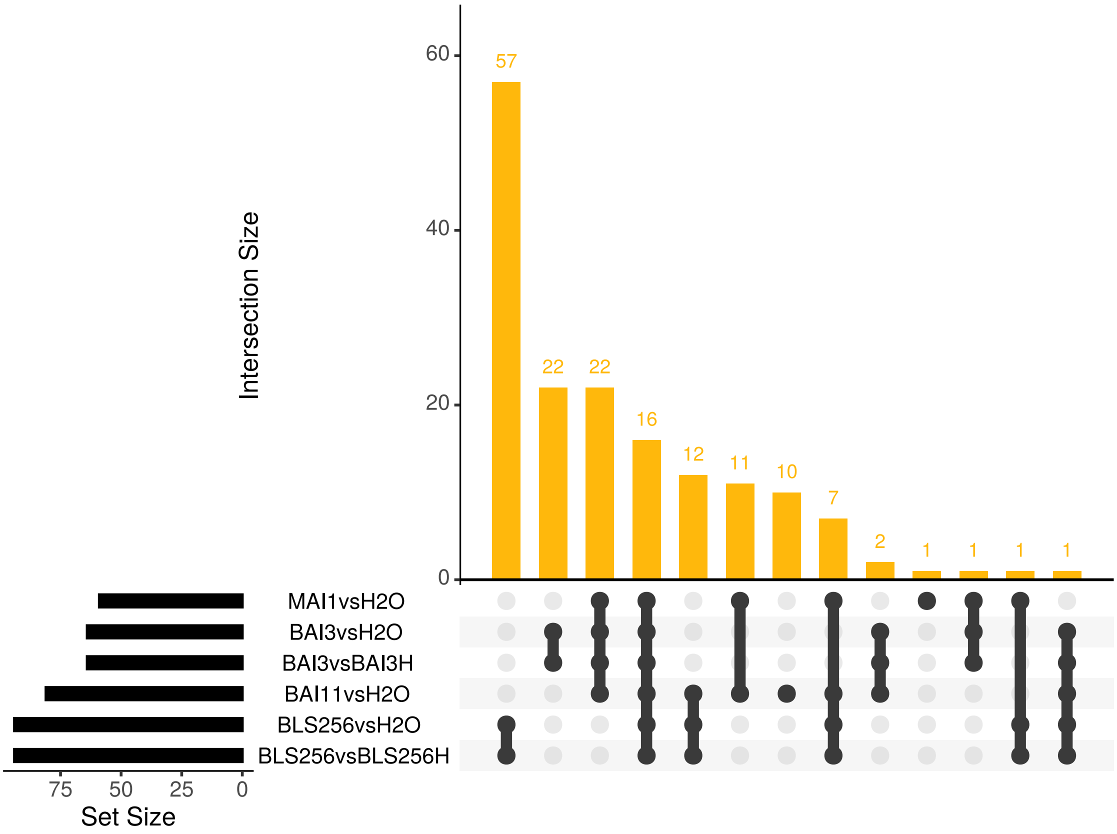

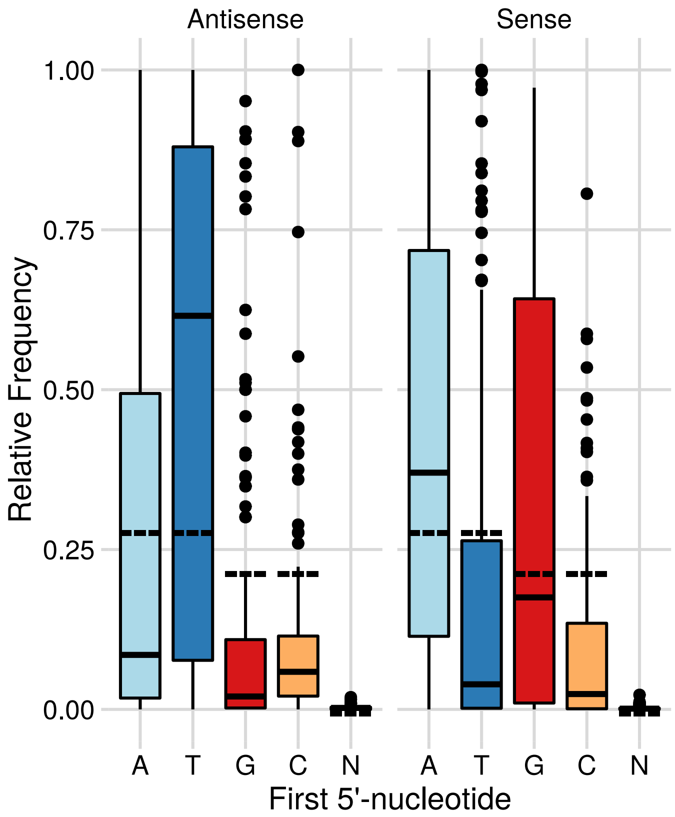

### Sup_Fig_S08.pdf

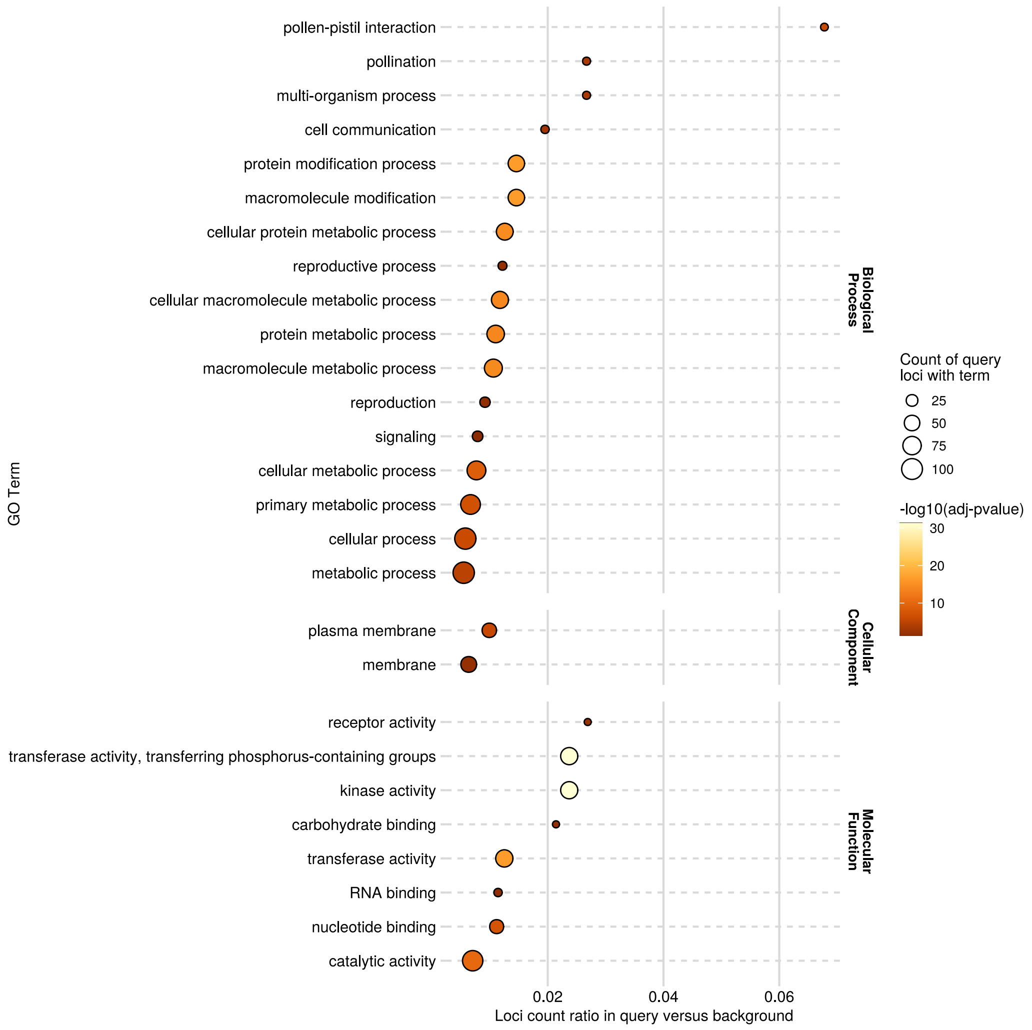

### Sup_Fig_S09.pdf

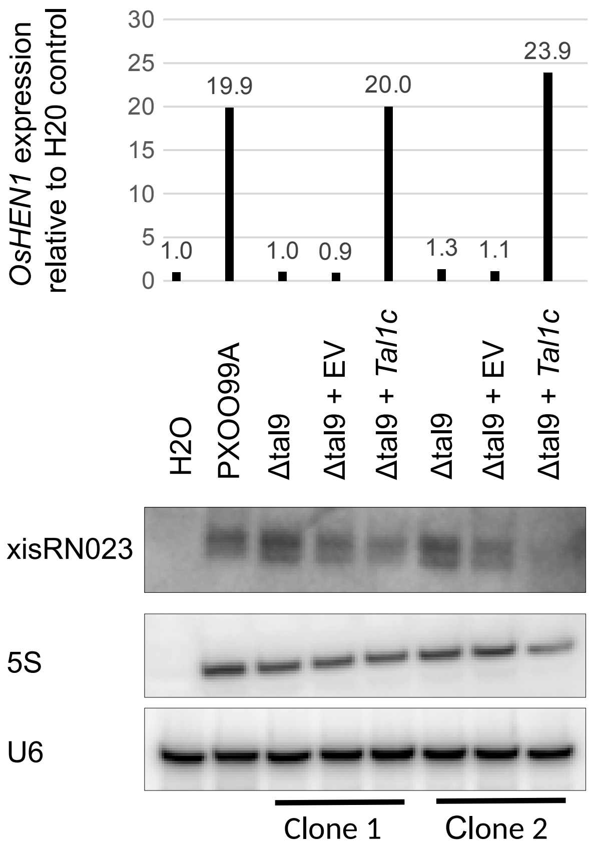

### Sup_Fig_S10.pdf

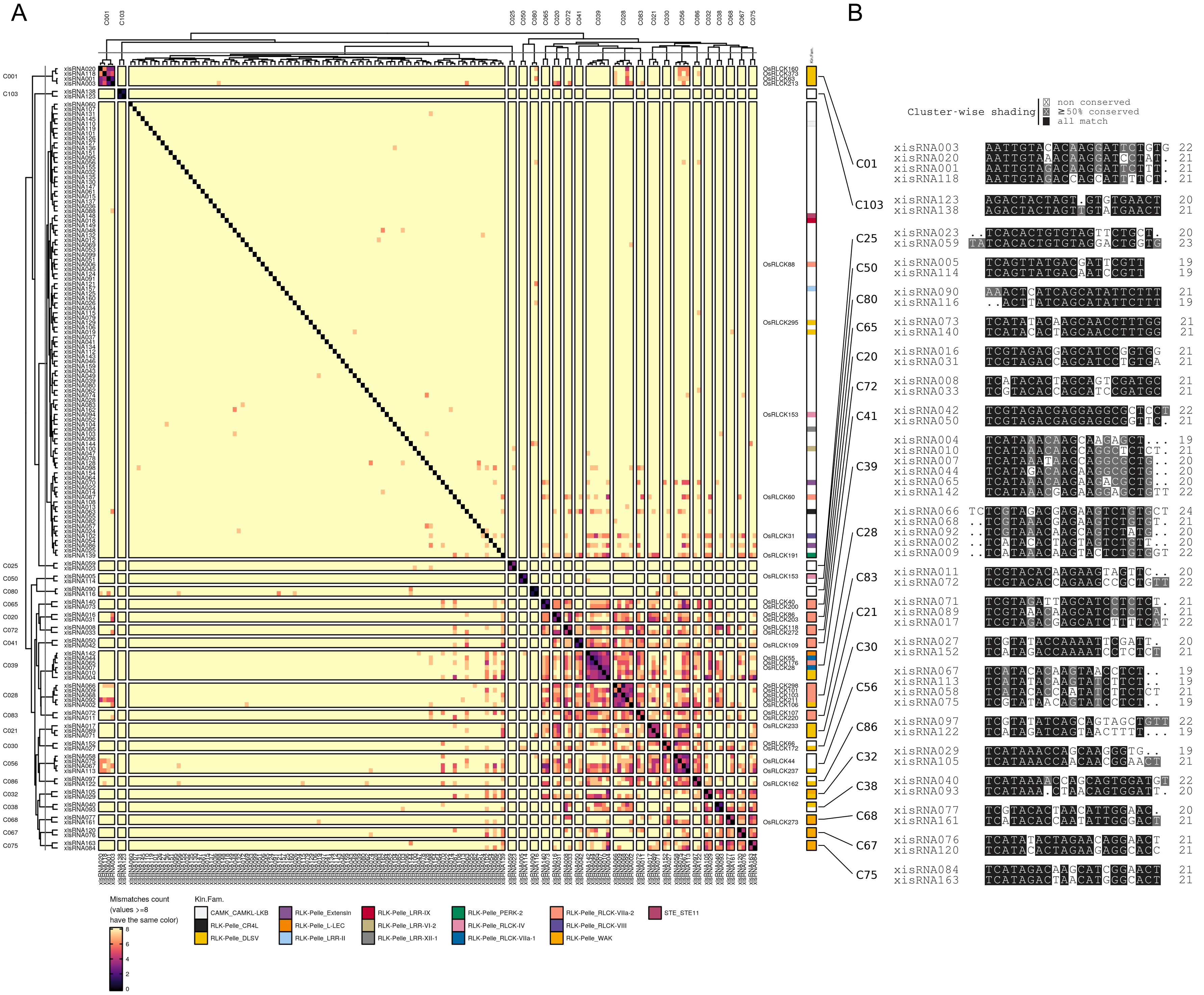

### Sup_Fig_S11.pdf

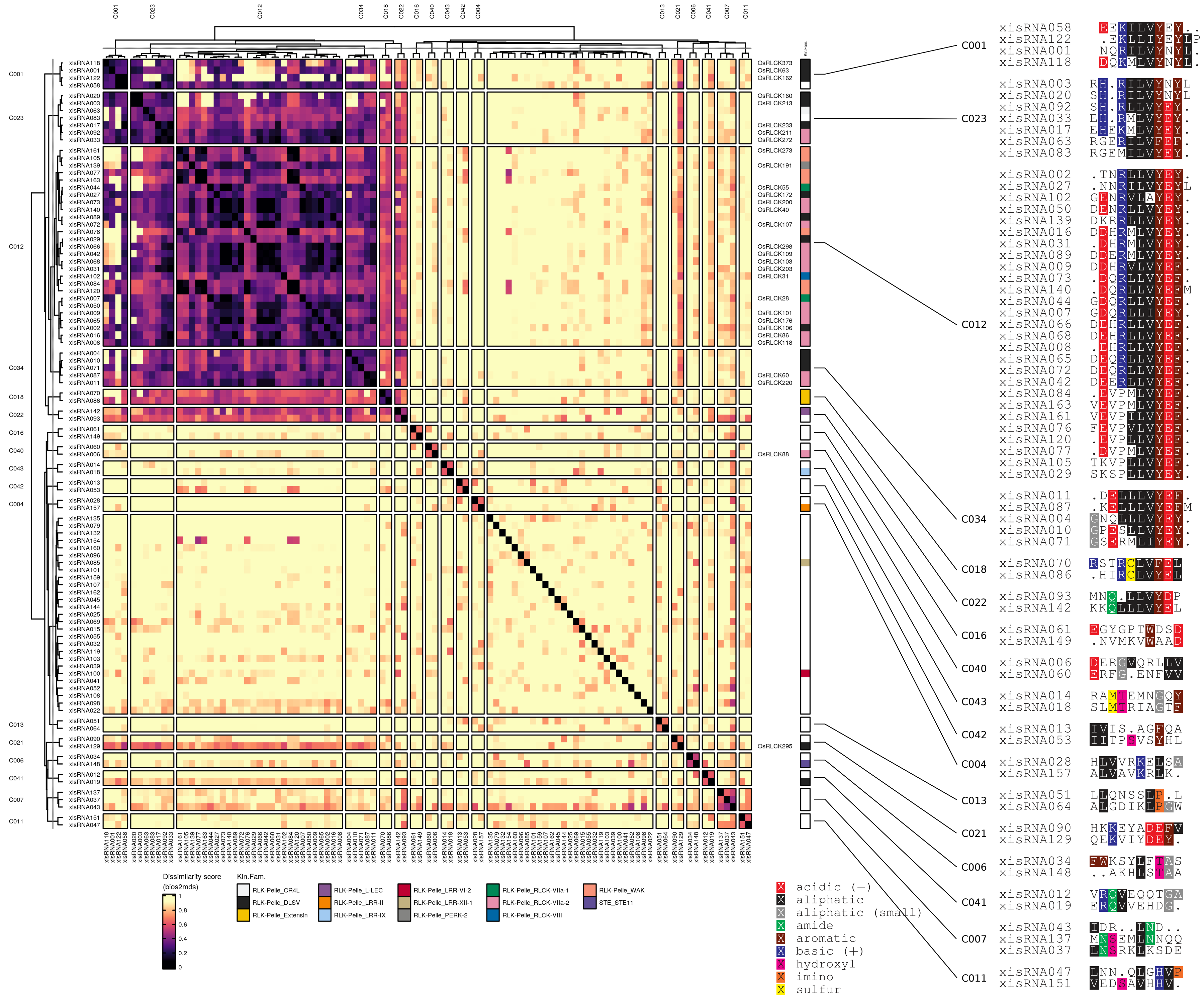

### Sup_Fig_S12.pdf

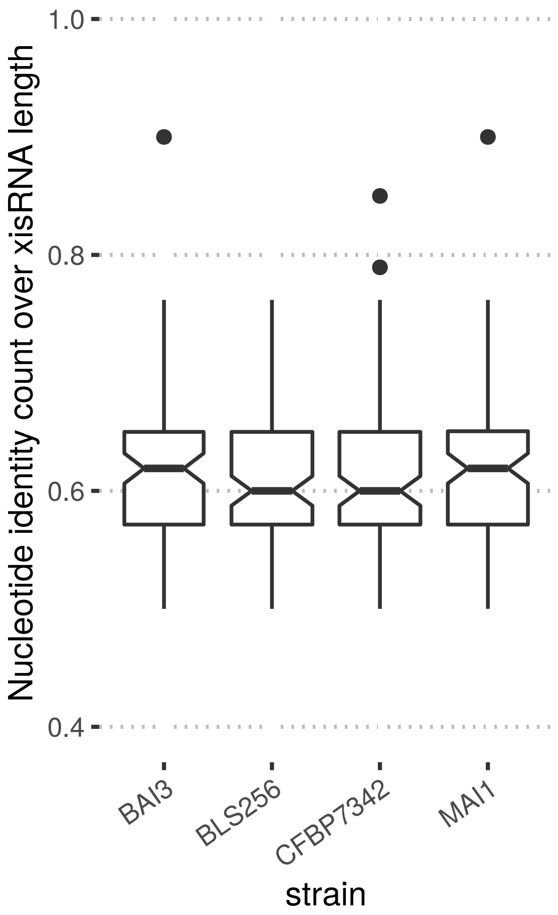
